# Quantifying protein-protein interactions by molecular counting with mass photometry

**DOI:** 10.1101/2020.01.31.925156

**Authors:** Fabian Soltermann, Eric D.B. Foley, Veronica Pagnoni, Martin R. Galpin, Justin L.P. Benesch, Philipp Kukura, Weston B. Struwe

## Abstract

Interactions between biomolecules control the processes of life in health, and their malfunction in disease, making their characterization and quantification essential. Immobilization- and label-free analytical techniques are particular desirable because of their simplicity and minimal invasiveness, but struggle to quantify tight interactions. Here, we show that we can accurately count, distinguish by molecular mass, and thereby reveal the relative abundances of different un-labelled biomolecules and their complexes in mixtures at the single-molecule level by mass photometry. These measurements enable us to quantify binding affinities over four orders of magnitude at equilibrium for both simple and complex stoichiometries within minutes, as well as to determine the associated kinetics. Our results introduce mass photometry as a rapid, simple and label-free method for studying sub-μM binding affinities, with potential to be extended towards a universal approach for characterising complex biomolecular interactions.

---

Understanding how biomolecules interact with each other is central to the life sciences. The complexity thereof ranges from specific binary interactions, such as between antibodies and antigens^[1–3]^, to the formation of complex macromolecular machines^[4,5]^. Conversely, undesired interactions are often associated with disease, such as the formation of protein aggregates in neurodegenerative disease^[6]^, or the engagement of a virus with its target cell^[7,8]^. The high-specificity and critical role of these interactions make them an ideal target for intervention, either in promoting a certain response by presenting an alternative binding partner, or preventing (dis)assembly^[9–11]^. This diversity comes with a broad range of binding strengths and dynamics, measured in terms of thermodynamic and kinetic quantities such as equilibrium constants (e.g. for dissociation, *K*_d_), free energies and rate constants (*k*_off_ and *k*_on_).

In broad terms, existing biophysical methods can be categorised into size-based approaches performing quantification and separation by size or diffusion coefficient, physical interaction with functionalised surfaces, direct mass measurement, enthalpy changes or light scattering^[12–17]^. These ensemble-based methods are complemented by fluorescence-based approaches^[18]^ capable of operating at the single molecule level, providing additional information on sample heterogeneity and dynamics ^[19,20]^. All of the above methods operate in the context of various practical shortcomings such as non-native environments, artefacts caused by protein immobilization and labelling, lack of sensitivity at low concentrations or lack of resolution^[21–23]^. Biological systems can pose additional challenges from particularly fast or slow kinetics to complexities arising from multiple co-existing species. Label-free methods struggle particularly for strong binding affinities (*K*_d_<μM), which are often encountered for interactions of particular relevance for biopharmaceuticals in the context of antibody-based drugs^[24]^.

We have recently developed mass photometry (MP), originally introduced as interferometric scattering mass spectrometry (iSCAMS), as a means for detecting and measuring the mass of single proteins and the complexes they form in solution. MP detects single biomolecules by their light scattering as they bind non-specifically to a microscope cover glass surface. Each binding event leads to a change in refractive index at the glass/water interface, which effectively alters the local reflectivity and can be detected with high accuracy by taking advantage of optimized interference between scattered and reflected light (**Figure 1a**)^[25]^. The magnitude of the reflectivity change can be converted into a molecular mass, for polypeptides with ~2% mass accuracy and up to 20 kDa mass resolution by calibration with biomolecules of known mass^[26]^. Both the original^[26]^, and subsequent studies have proposed methods to extract binding affinities from MP distributions of biomolecular mixtures^[27]^, and shown that the obtained affinities agree broadly with alternative approaches ^[26–28]^. The degree to which these MP distributions are indeed quantitative, and how they can be used to efficiently extract not only binding affinities but also kinetics, however, remain unexplored.

**Figure 1.**
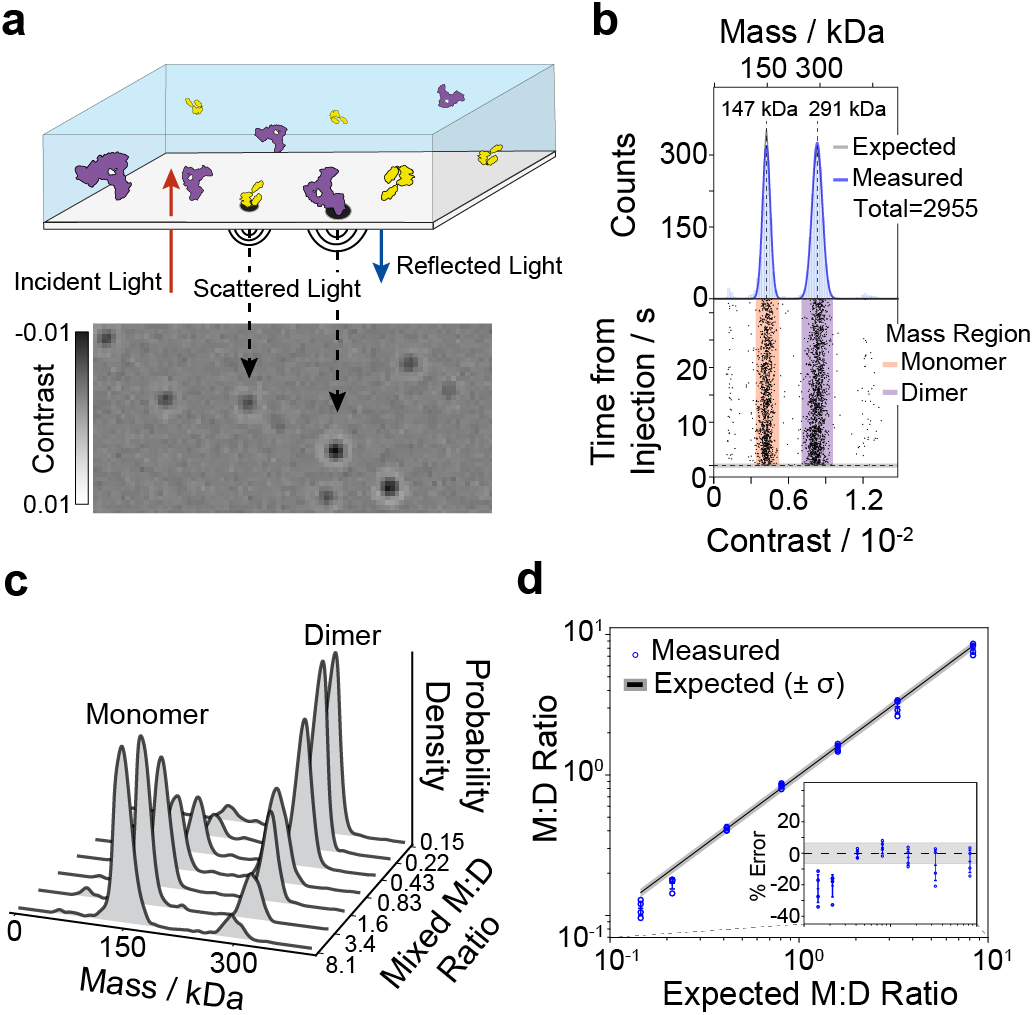
Principle of single molecule counting by mass photometry. (**a**) Label-free single molecule detection by imaging the interference of scattered and reflected light arising from individual protein landing events over time. (**b**) Scatter plot of single molecule contrasts and resulting mass distribution for a 1:1 monomer:dimer 2G12 mixture. (**c**) Mass distributions for varying 2G12 monomer:dimer ratios. (**d**) Comparison of monomer:dimer ratios measured by MP compared to expectations based on UV-VIS absorption characterisation.

Label-free single molecule detection in principle provides the purest and most direct measurement of sample concentration by counting individual molecules. To explore this capability in the context of biomolecules, we chose monomers and dimers of the HIV-1 neutralizing antibody 2G12 (**Supplementary Figures 1-3**), which produced mass distributions with the expected major bands at 147 kDa and 291 kDa (**Figure 1b**). Repeating these experiments for monomer/dimer ratios ranging from 0.15 to 8.1 (**Figure 1c**) revealed close agreement with UV-VIS-based characterisation within the experimental error (4.6% RMS), except for noticeable deviations (~20%) for the lowest ratios (**Figure 1d**). We found that such deviations could almost exclusively be attributed to sample preparation, such as an additional dilution step for the data shown (**Supplementary Figure 4 & 5**).

Equipped with these benchmarking results, we set out to investigate the suitability of MP to characterise interactions of varying affinities, using the immunoglobulin G (IgG) monoclonal antibody trastuzumab (Herceptin®) binding to soluble domains of IgG Fc receptors or ErbB2 (HER2) antigens. Trastuzumab, herein referred to as IgG, and Fc*γ*RIa by themselves revealed monodisperse distributions at 154 ± 1 kDa and 50 ± 1 kDa, respectively (**Figure 2a, Supplementary Figure 6**). A 1:1 Fc*γ*RIa-IgG mixture resulted in a large IgG-Fc*γ*RIa complex peak, corresponding to ~90% complex formation, from which we can extract an apparent *K*_d_ = 50 ± 10 pM by counting bound and unbound species in combination with knowledge of the total protein concentration (more information in **Supplementary Equations 1-6, Supplementary Figure 7**). IgG N-glycan removal (**Supplementary Figure 8**) weakened FcR binding^[29]^ resulting in a 1:1 mixture of Fc*γ*RIa and deglycosyated IgG exhibiting considerably less bound antibody (~50%) (**Figure 2b**), corresponding to an apparent *K*_d_ = 1.0 ± 0.1 nM (**Supplementary Figure 9**).

**Figure 2.**
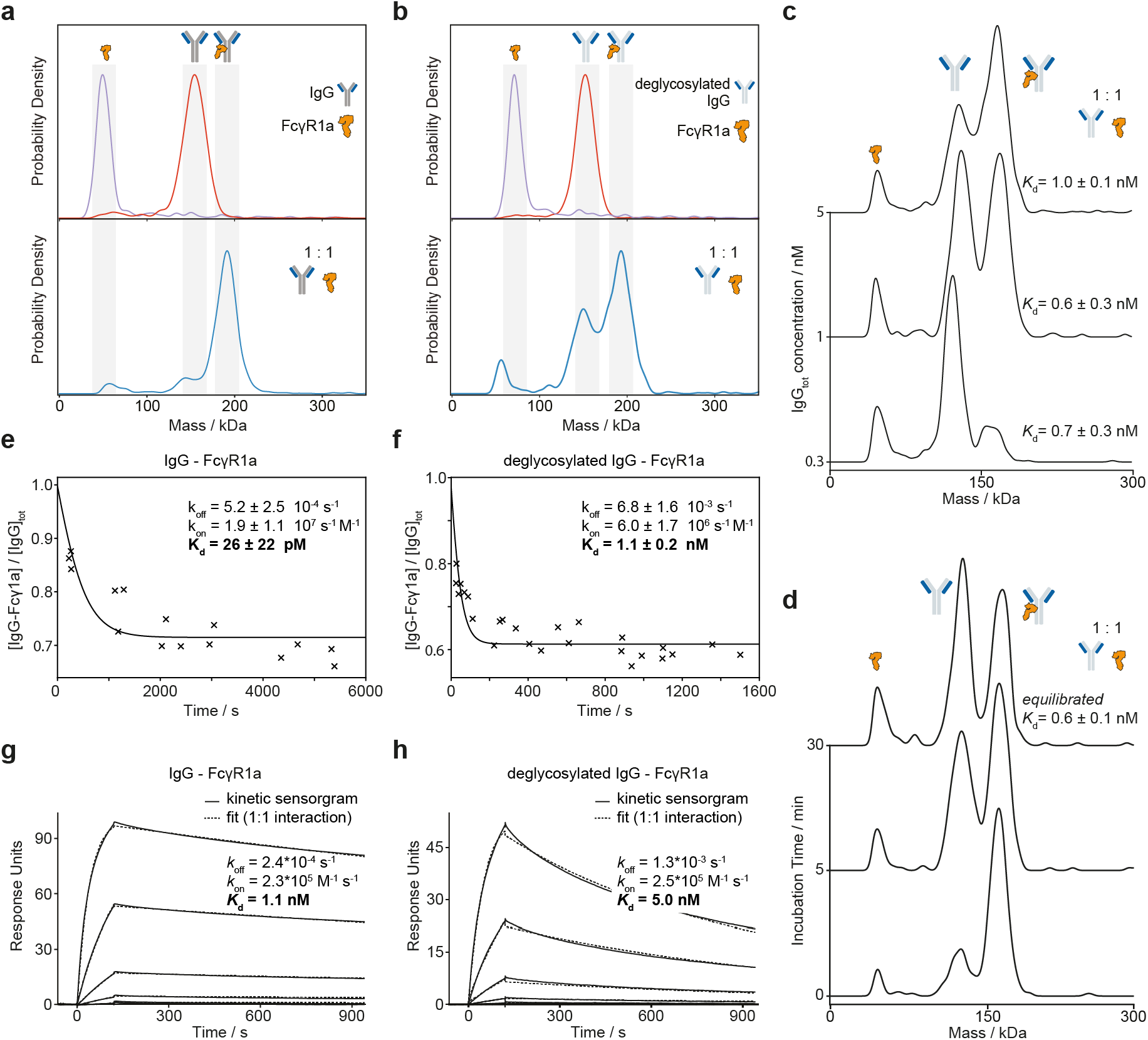
Single-shot *K*_d_ and kinetics of IgG-FcγRIa interactions. (**a**) MP mass distributions of IgG (red), FcγRIa (purple) and a 1:1 mixture of IgG-FcγRIa (blue). (**b**) MP distributions of deglycosylated IgG (red), FcγRIa (black) and 1:1 mixture of IgG-FcγRIa (blue). (**c**) Mass distributions for a 1:1 mixture of deglycosylated IgG-FcγRIa at total IgG concentrations ranging from 300 pM to 5 nM and respective *K*_d_ calculated from a single-shot measurement. (**d**) Mass distributions for a 1:1 mixture of deglycosylated IgG-FcγRIa at 1.5 nM total IgG concentration, ranging from 0 to 30 minutes after dilution from 2.9 μM. (**e,f**) Mole fraction of assembled IgG-FcγRIa and deglycosylated IgG-FcγRIa complexes as a function of time after dilution from 2.7 μM to 0.3 nM total IgG concentration, and 2.6 μM to 5 nM total deglycosylated IgG concentration and corresponding single exponential fits. **(g,h)** Corresponding SPR analysis of IgG-FcγRIa and deglycosylated IgG-FcγRIa (**h**).

This simple single-shot approach presented so far, however, necessarily neglects the importance of kinetics and equilibration conditions. To address this, we probed Fc*γ*RIa binding to deglycosylated IgG at a 1:1 ratio and 5, 1 and 0.3 nM final protein concentrations (**Figure 2c**, **Supplementary Figure 10**). At concentrations above the *K*_d_ we found mostly bound complexes, with free species dominating below the *K*_d_, but all measurements providing similar binding affinities (*K*_d_ =1.0 ± 0.1, 0.6 ± 0.1 and 0.7 ± 0.3 nM), suggesting that they were performed at or close to equilibrium (**Supplementary Figure 11**). These binding affinities were confirmed after equilibration time screening (**Supplementary Figures 12 & 13**).

For quantification of the expected tighter interaction between Fc*γ*RIa and IgG, the screening at a range of concentrations was essential to ensure the observed mass distributions were representative of the interaction to be quantified (**Supplementary Figures 14-16**). As an additional example, for the HER2-IgG interaction, a simple single-shot experiment at nM concentration would have led to a *K*_d,1_ = 1.4 ± 0.1 nM and *K*_d,2_= 4.8 ± 0.3 nM (**Supplementary Figure 17**). By simply recording distributions at a few different concentrations, however, we could reveal a linear dependence of our K_d_ values on concentration, indicating a very tight K_d_ < 70 pM, and/or slow interactions with off-rates on the order of hours. Therefore, performing a few measurements at a range of concentrations is crucial to prevent misinterpreting data derived from a single-shot *K*_d_ approach for very strong interactions. Irrespective, our approach provides a rapid and clear distinction between interactions with vastly different binding affinities, which only need to be refined if highly accurate measurements are required.

The importance of (dis)association rates in addition to thermodynamic quantitites raises the question to which degree we can use MP to directly visualise and quantify interaction kinetics. As MP measurements currently take place in the <100 nM concentration range, we should be able to access disassociation kinetics by simply diluting to total protein concentrations around the estimated *K*_d_ (approx. 1:1 ratio bound:unbound species for a 1:1 interaction), and monitoring the bound:unbound ratio throughout (**Figure 2f, Supplementary Figure 18a**). The observed exponential decay of a complex upon rapid dilution below the *K*_d_ reveals the desired kinetic information, while the plateau yields the *K*_d_, ultimately enabling us to also determine *k*_off_ and *k*_on_. For Fc*γ*RIa binding to deglycosylated IgG, this approach yielded *K*_d_ = 1.1 ± 0.2 nM in good agreement with our single shot measurements (**Figure 2c**), *k*_on_ = 6.0 ± 1.7 × 10^6^ M^−1^ s^−1^, and *k*_off_ = 6.8 ± 1.6 × 10^−3^ s^−1^. The corresponding experiment with glycosylated IgG-Fc*γ*RIa yielded *K*_d_ = 26 ± 22 pM with an off-rate one order of magnitude slower (5.2 ± 2.5 × 10^−4^ s^−1^) than for deglycosylated IgG but an almost identical on-rate (1.9 ± 1.1 × 10^7^ M^−1^ s^−1^) (**Figure 2e**), again in good agreement with our single-shot screening data (**Supplementary Figures 15 & 16**). The difference in *K*_d_ between glycosylated and deglycosylated IgG originates mostly from the off-rate demonstrated by protein-protein interactions (**Supplementary Figure 19**), confirming that the glycans are critical in determining the binding strength. Association measurements (**Supplementary Figure 18b, 20 & 21**) can in principle be used in an analogous fashion, although we found them more susceptible to protein loss due to non-specific adsorption (**Supplementary Figures 22–24**). Overall, our results were in good agreement with SPR measurements (**Figures 2e & 2f**), subject to on-rate differences expected between a matrix and surface-immobilization based approach compared to ours, where all interactions take place in free solution (**Supplementary Figures 25**).

A key advantage of MP over ensemble-based approaches is our ability to easily distinguish between different species contributing to a multicomponent system, as given by the IgG:FcRn interaction involving as many as five different interacting species. FcRn regulates serum IgG half-life and transcytosis to the fetus via a pH gradient in endosomes, yet the interplay between self-assembly and IgG binding is disputed^[30,31]^, which are both important factors in biotherapeutic design. Based on the existing literature we based our calculations on the independent free monomer binding model (**Supplementary Figure 26a)**. At pH = 5, FcRn formed monomers and dimers with a *K*_d_ = 31 ± 11 nM (**Figure 3a and Supplementary Figure 27**). At pH = 5.5 and pH = 6, only negligible amounts of FcRn dimers were present with a FcRn monomer-dimer *K*_d_ >200 nM (**Supplemental Figures 28 & 29**). The resulting *K*_d_ values for the IgG-FcRn interaction, at pH 5, were 44 ± 9 nM for the monomer-dimer equilibrium, 59 ± 8 nM for the IgG-FcRn_monomer_ and 6.6 ± 0.6 nM for IgG plus two FcRns (**Supplementary Figures 30 and Supplemental Equations 7-18**). Increasing the pH to 5.5 decreased the binding affinities to 171 ± 19 nM and 225 ± 20 nM but did not significantly affect the binding affinity of IgG plus two FcRns of 3.9 ± 1.5 nM (**Figure 3b, Supplementary Figure 31 & 32**), contrasting SPR results for similar systems reporting an ensemble *K*_d_ =760 ± 60 nM for all interactions^[31]^. At pH = 6 and 7, our current sensitivity only allowed for an estimate of the binding affinities to be >200 nM (**Figure 3b, Supplementary Figures 33 and 34**). These results highlight pH-dependent FcRn dynamics and IgG engagement, and reveals cooperativity where the second receptor binds IgG tighter than the first and with a weaker pH sensitivity.

**Figure 3.**
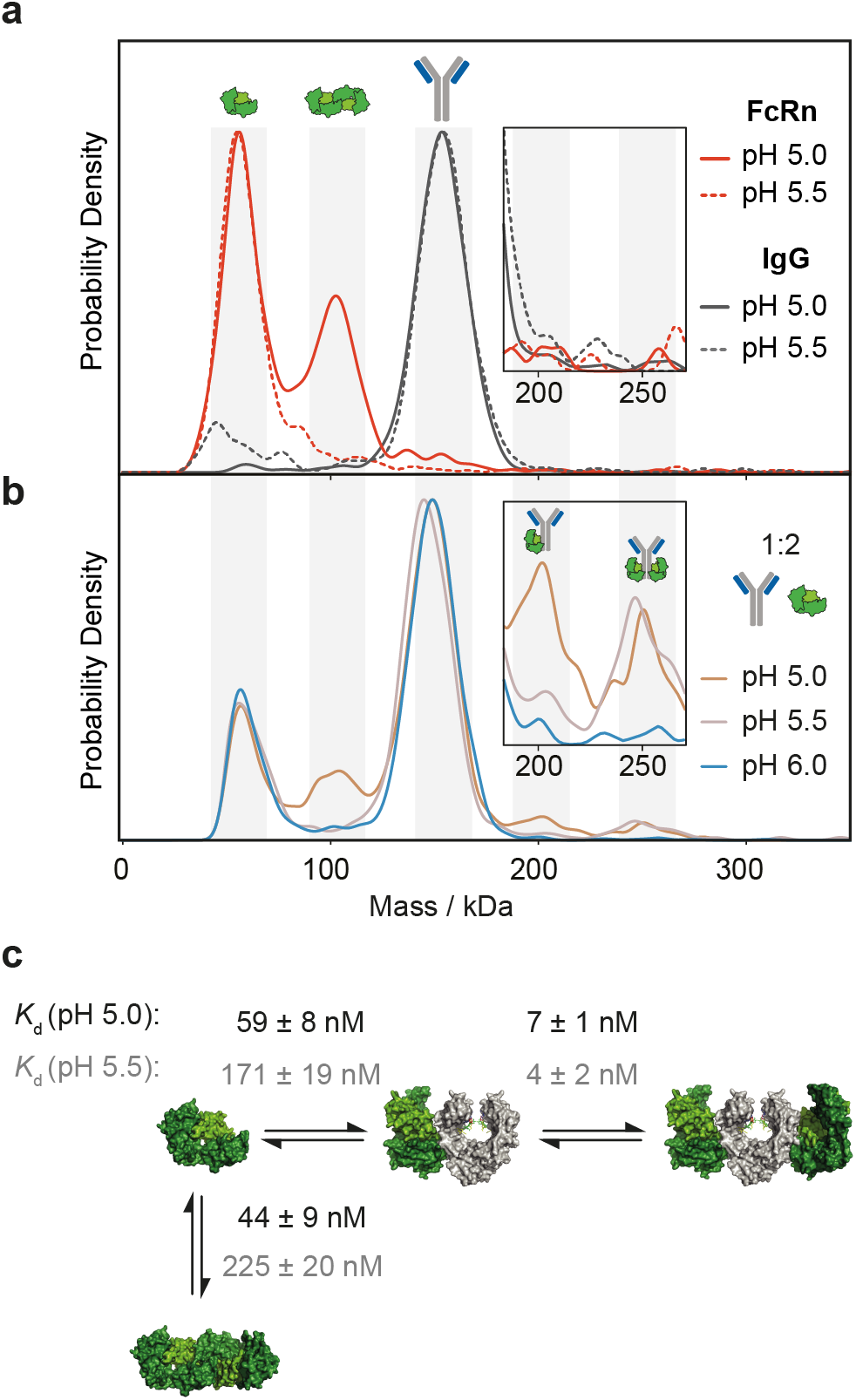
Binding stoichiometry and affinity of the IgG-FcRn interaction as a function of pH. (**a**) Self-assembly of FcRn dimers at pH 5 (red) and 5.5 (dotted red) and equivalent pH measurements of IgG at pH 5 (grey) and 5.5 (dotted grey). (**b**) IgG-FcRn complexes (1:1 mixture) at pH 5, 5.5 and 6. (**c**) Associated pH dependent binding affinities of interaction revealing cooperativity in FcRn binding.

Taken together, we have demonstrated that molecular counting with MP is sensitive, quantitative and accurate in determining the relative abundances of different biomolecules and their complexes in solution. When implemented in the vicinity of the binding affinity, a single measurement lasting typically 30 s, or 240 s for continuous flow injection, yields accurate binding affinities spanning four orders of magnitude from 30 pM to 200 nM, while enabling kinetic probing with a time-resolution on the order of 30 s in the range of minutes to hours. As a result, MP affords real-time assessment of (dis)assembly completely label-free and entirely independent of immobilization, minimising any possible perturbations, as well as being intrinsically sensitive to binding stoichiometries and oligomerization. The current limitation to sub-μM affinities and concentration range can be addressed in the future by a combination of surface passivation approaches^[32]^, as well as improvements to hardware and software, with which we expect to reach the μM range in the near future. This will enable measurements up to 100 μM affinities, making MP a powerful approach for characterising biomolecular interactions without labels and single-molecule sensitivity in a minimally perturbative fashion. Furthermore, the applicability of MP to both nucleic acids^[33]^ and large multimolecular machines provides scope for MP becoming a universal tool for studying biomolecular interactions and dynamics in a rapid, label-free and yet single molecule sensitive fashion.

## Abbreviations

MP: mass photometry
iSCAMS: interferometric scattering mass spectrometry
SPR: surface plasmon resonance
IgG: immunoglobulin G
FcγR: Fc-gamma receptor
FcRn: neonatal Fc receptor
*K*_d_: dissociation constant

## Acknowledgments

We thank Mohammed Khan and Ben Davis, University of Oxford, for supplying deglycosylated trastuzumab and discussions, Stephan Burkhalter, ETH Zurich and PSI, for contributions to the analysis code, Andrew Baldwin, University of Oxford for helpful discussion, David Staunton, University of Oxford for assisting in SPR measurements, and Daniel Cole for the construction of the mass photometer. This work was supported by funding from the Engineering and Physical Sciences Research Council (EPSRC) and Medical Research Council (MRC) [grant number EP/L016052/1]. The authors recognize support from an ERC Consolidator Grant (PHOTOMASS 819593).

## Author contributions

Conceptualization: F.S., W.S., and P.K.; Investigation: F.S., E.F. and V.P.; Formal Analysis: F.S. and M.G.; Writing – Original Draft: F.S., W.S., and PK; Writing – Review & Editing: all authors; Visualization: F.S., W.S. and J.L.P.B.; Supervision: W.S. and P.K.

## Materials & Methods

We first characterised all compounds separately by MP to assess their oligomeric composition (**Supplementary Figure 10**). Subsequently, we prepared stoichiometry-dependent mixtures of analyte and binding partner (1:1 or 1:2) at μM concentration and equilibrated at room temperature overnight. The samples were then diluted to concentrations near the expected *K*_d_ value and equilibrated during a time *t* before conducting a concentration screening (**Supplementary Figure 10**). In these experiments we set the equilibration time to *t* = 15 min and measured the sample at three different dilutions (0.3 nM, 1 nM, 5-50 nM) to obtain an estimate of the *K*_d_ and, most importantly, to confirm that complexes are dissociating upon dilution, i.e. the signal corresponding to the biomolecular complex decreases in relative intensity (see **Figure 2c & 2d**). If complexes do not dissociate upon dilution, i.e. relative peak intensities are not changing, the biomolecule interaction is likely to exhibit very tight binding affinities (*K*_d_ < 0.1 nM) and/or slow kinetics (> 5 hrs) (**Supplementary Figure 15**) possibly beyond the dynamic range and sensitivity of the current MP instrumentation (**Supplementary Figure 17)**. Provided the *K*_d_ screening was successful we proceeded with the kinetic studies for which we diluted the sample to a concentration around the estimated *K*_d_. After dilution/mixing we followed dissociation/association of the biomolecular complex by acquiring individual MP measurements at different time points. From these experiments, we obtained *K*_d_ values and on-and off-rates. With the current dynamic range we can inject concentrations from 100 pM to 50 nM, and thus determine quantitate binding affinities ranging from 10 pM to 300 nM and measure kinetics ranging from a few minutes to several hours.

The 2G12 antibody was expressed and purified as described previously^[34]^. Briefly, Plasmids encoding 2G12 antibody heavy and light chains were transiently expressed in HEK 293F at a cell density of 1 × 10^6^ cells/mL with a 1:2 construct ratio (heavy to light chain). After 5 days, cells were pelleted and the supernatant was loaded onto a HiTrap Protein A column (GE Healthcare) following the manufacturers protocol. The binding buffer was 20 mM sodium phosphate, pH 7.5, and the elution buffer was 0.1 M citric acid, pH 3. Antibodies were immediately neutralized with 1 M Tris-HCl, pH 9, prior to buffer exchange into phosphate buffered saline (PBS) (Gibco® DPBS) using a 50 kDa cut-off spin filter (Vivaspin, Sartorius). Antibody oligomers were then separated with size exclusion chromatography (HiLoad Superdex 16/600 200 ps, GE Healthcare) and fractions were collected according to the SEC profile and SDS-PAGE (NuPAGE™ 4-12% Bis-Tris Protein Gels, Invitrogen™). Antibodies were stored at 4°C. Trastuzumab (Herceptin®) was purchased from Oxford University Hospital, CD64 human recombinant (His Tag) was purchased from SinoBiological, recombinant human FcRN (His and Avi tagged) was purchased from Stratech Scientific Ltd, ErbB2 (Her2) human recombinant was purchased from Sigma-Aldrich. Deglycosylated trastuzumab was obtained by treatment with Endo S (New England BioLabs) at 37 °C, overnight. 2G12 antibody was used for contrast vs. molecular weight calibration, with monomer and dimer masses of 148.5 and 296.9 kDa respectively (**Supplementary Figure 3**).

### Sample Preparation

Samples and buffers were filtered through 0.1 μm centrifugal filters (Ultrafree -MC- VV, MerckMillipore) prior to use. Initial stock concentrations of all proteins were measured using a DeNovix DS-11+ spectrophotometer. Protein solutions were diluted in PBS (Gibco DPBS (1X)) to obtain concentrations between 0.2 to 1 absorbance at 280 nm (low μM range). Pipetting precision was checked with a microbalance (Mettler Toledo, AT261).

#### Relative abundance measurements

For the experiments determining the accuracy of determining relative abundances (**Figure 1a-d**), we used the two SEC-fractions of the 2G12 antibody purification (**Supplementary Figure 1**), containing either the pure monomer or the intermolecular domain exchanged dimer version of the monomer^[35]^. Characterizion of the monomer fraction with MP confirmed the absence of dimerization and analogously the dimer fraction revealed absence of dissociation into monomer. Different ratios of the two non-interacting species (monomer and dimer) were mixed at μM concentration and diluted in PBS to pM-nM concentrations prior to measurement (**Table 1**). For the two lowest M:D ratios (0.15 and 0.22) in **Figure 1c & 1d** the pure monomer stock solution had to undergo an additional step from 1 μM to 500 nM. All pipetting steps were checked on a microbalance.

**Table 1:**
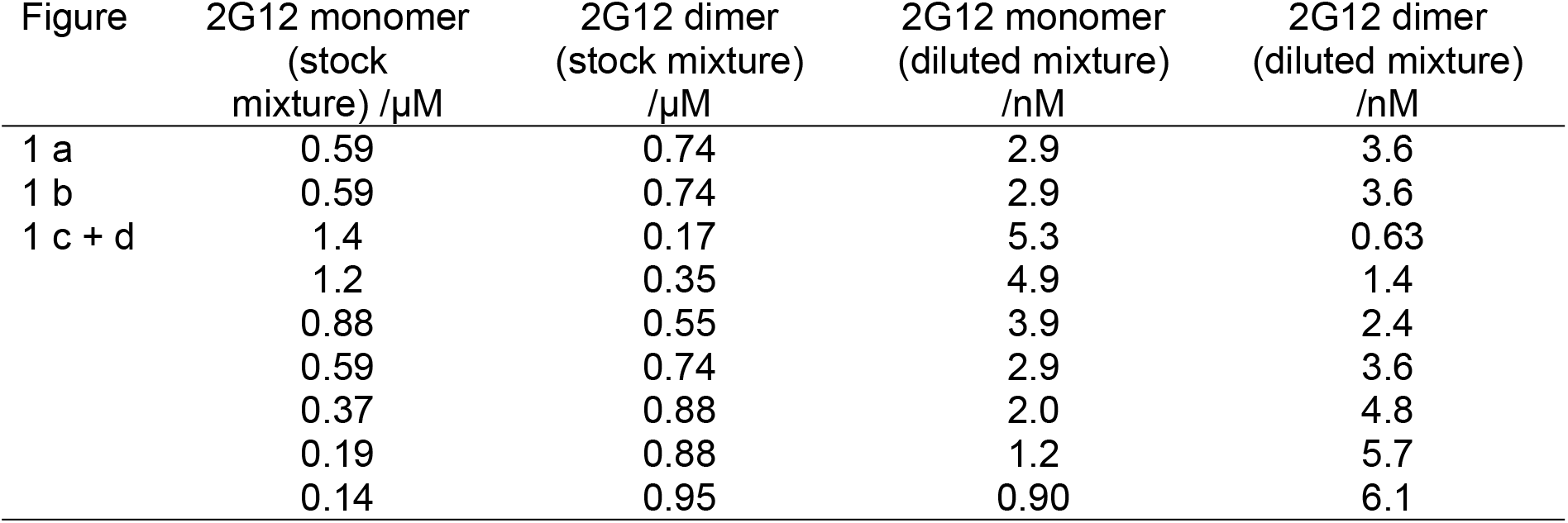
2G12 stock solutions and diluted concentrations.

| Figure | 2G12 monomer<br>(stock<br>mixture) /μM | 2G12 dimer<br>(stock mixture)<br>/μM | 2G12 monomer<br>(diluted mixture)<br>/nM | 2G12 dimer<br>(diluted mixture)<br>/nM |
| --- | --- | --- | --- | --- |
| 1 a | 0.59 | 0.74 | 2.9 | 3.6 |
| 1 b | 0.59 | 0.74 | 2.9 | 3.6 |
| 1 c + d | 1.4 | 0.17 | 5.3 | 0.63 |
|  | 1.2 | 0.35 | 4.9 | 1.4 |
|  | 0.88 | 0.55 | 3.9 | 2.4 |
|  | 0.59 | 0.74 | 2.9 | 3.6 |
|  | 0.37 | 0.88 | 2.0 | 4.8 |
|  | 0.19 | 0.88 | 1.2 | 5.7 |
|  | 0.14 | 0.95 | 0.90 | 6.1 |

#### *Screening experiments* (Figure 2a-d and Figure 3)

Individual FcRs (Fc*γ*RIa, FcRn) and IgGs (trastuzumab and deglycosylated trastuzumab) were mixed at μM concentration in a 1:1 (IgG-Fc*γ*RIa, deglycosylated IgG-Fc*γ*RIa) or 1:2 (IgG-FcRn) ratio and incubated overnight at room temperature (see Table 2 for exact concentrations). The μM mixtures were diluted in PBS to pM-nM concentrations and measured at specific incubation times (Table 2). For FcRn-IgG, this procedure was repeated for each pH (5.0, 5.5, 6.0, 7.2) (Figure 3). *Kinetic experiments* (Figure 2e & 2f): Dissociation experiments were prepared as described for those shown in Figure 2a-f and repeated after specific incubation times. For association experiments, individual proteins were diluted in PBS to pM-nM concentrations and then mixed in 1:1 ratios. Measurements were taken after specific incubation times had elapsed. For association experiments of IgG and Fc*γ*RIa (shown in Supplementary Figure 20 & 21), the pure compounds were diluted to nM concentrations, mixed, then incubated for the desired incubation time and then measured by MP. The sample tube passivation protocol, based on Casein was applied (see below).

**Table 2:** FcR/IgG mixtures and incubation times

| | Figure | FcR<br>(stock) /μM | IgG<br>stock)<br>/μM | FcR<br>(diluted<br>mixture)<br>/nM) | IgG<br>(diluted<br>mixture)<br>/nM | incubation<br>time after<br>dilution<br>/min |
| --- | --- | --- | --- | --- | --- | --- |
| screening<br>experiments | 2 a | 2.7 | 2.7 | 4.4 | 4.4 | 10-18 |
|  | 2 b | 2.9 | 2.9 | 4.6 | 4.6 | 12-18 |
|  | 2 c | 2.9 | 2.9 | 0.3, 1.3, 4.6 | 0.3, 1.3, 4.6 | 10-35 |
|  | 2 d | 2.9 | 1.3 | 1.3 | 1.3 | 0.4-30 |
|  | 3a | 18.8 | 7 | 11.8 | 9.2 | 0.4-89 |
|  | 3b | 16.9 | 7.6 | 0.4-6.0 | 0.2- 2.7 | 0.5-78 |
| kinetic<br>experiments | 2e | 2.7 | 2.7 | 0.21- 0.26 | 0.21- 0.26 | 1.8-88 |
|  | 2f | 2.6 | 2.6 | 4.6- 4.9 | 4.5- 4.8 | 0.3-58 |

#### ErbB2-trastuzumab interaction study

Experiments were performed analogously to IgG-FcγRIA. Briefly, we applied the *K*_d_ screening procedure to investigate the binding affinity of trastuzumab to the ErbB2 (HER2) antigen. The two compounds were mixed in a 1:1 molar ratio and incubated at room temperature overnight. The next day, MP data was acquired at different dilutions (0.3, 1.3, 4.6 nM trastuzumab concentration) and time points after dilution (2.5 – 100 min). Gaskets (see below) were used for the experiments with the 1.3 and 4.6 nM dilutions and the continuous flow setup for the 0.3 nM dilution (Supplementary Figure 4).

### Mass Photometry

Microscope coverslip cleaning and assembly was performed as described previously^[26]^. Borosilicate microscope coverslips (No. 1.5H thickness, 24 × 50 mm, VWR (630-2603) were cleaned by 5 min sequential sonication (PS-40A Ultrasonic Cleaner, Cole Palmer) in Milli-Q water, isopropanol and Milli-Q water, followed by drying under a clean stream of nitrogen. Quality control of new batches of coverslips was based on checking the surface roughness on the mass photometry native images for 6 out of 100 coverslips from a single box. Batches with large particles or irregularities on the glass surface were disposed. Clean coverslips were assembled for sample delivery using silicone gaskets (CultureWell™ reusable gasket, 3mm diameter × 1 mm depth, Grace Bio-Labs) or plastic flow chambers (sticky-Slide VI 0.4, Ibidi). For experiments involving flow chambers, a microfluidic syringe pump-based system was used (World Precision Instruments AL-1000) (Supplementary Figure 4).

For data acquisition, gaskets were filled with buffer (40 μl) to enable focusing of the glass surface. Subsequently, 36 μl were removed, while keeping the focus position stable, and replaced by 36 μl sample solution at nM concentration. Measurements were started immediately (delay time ~2 seconds) for 30 - 60 seconds. For samples with sub-nM *K*_d_ values, we conducted continuous flow experiments at sub-nM concentrations typically over 240 s. Flow chambers were filled with 150 μl buffer and connected to the sample reservoir and syringe pump, while preventing the generation of air bubbles in the flow chamber. The flow chamber was then placed on the instrument in a way that the laser was positioned close to the inlet of the flow chamber. Next, the syringe pump was started in pump mode and run at 90 μl /min. Once the sample reservoir was almost empty, the sample (sub-nM concentration) was injected, focus checked and the measurement started. The acquisition time was chosen depending on concentration, i.e. lowering the concentration by a factor of 10 requires 10 times longer acquisition times to keep the total number of molecular counts constant. Continuous flow experiments at concentrations in the nM range or higher should be avoided, because the coverslip glass surface becomes saturated with analyte molecules within 1 min after injection.

The experimental mass photometry setup has been described elsewhere ^[25,26]^. Briefly, a 525 nm laser diode was used for illumination with the following instrument parameters: acquisition camera frame rate = 955 Hz, pixel binning = 4×4, 5-fold time-averaging. Data was acquired for 30 or 240 s, depending on the experiment. Particle detection and quantification was processed as described by Young *et al*. using custom software written in Python^[26]^. For each particle a time stamp, position and contrast value was obtained.

#### Quality Control for Quantitative Measurements

During acquisition each video was examined visually and excluded when one of the criteria listed in Table 3 was met. During data analysis we applied an xy-distance constraint filter to all confirmed counts, excluding particles which were recorded at the same xy-position in the field-of-view. This helped to exclude “blinkers” (i.e. fast binding and unbinding events at the same position) and significantly lowered the baseline/noise. For gasket experiments, we plotted the counts as a detection rate (−Δcounts/Δt) of the measured species X vs. time. The resulting decay curve was exected to follow a single exponential decay, which indicates an error-free experiment. We experienced deviations from a smooth decay when one or several of the criteria in Table 3 were given.

**Table 3:** Quality Data Criteria & Indicators (i.e. low molecular weight resolution and high noise level)

| Criteria | Indicators | Most common reasons |
| --- | --- | --- |
| Out-of-focus measurement | Inversed contrast or anomalous PSF of particles in ratiometric image or no counts | Wrong focus position |
| Unstable focus | Focus voltage is drifting | Excessive immersion oil between stage and coverslip |
| Rough/irregular glass surface | Large interfering black dots in native image | Insufficient quality control during glass cleaning procedure |
| Unbinding events | Particles with negative contrast values | Unsuitable pH, unsuitable glass surface properties (test APTES coating)<sup>[37]</sup> |
| Buffer impurities | Large particles floating around and causing interferences in the ratiometric image | Buffer or sample not filtered suitably (i.e. 0.22 µm filter), sample is precipitating under chosen conditions |
| Too high concentration | Particles are overlapping in time and position | Incorrect dilution |

### Data Analysis

#### Quantification of species-specific counts

Contrast values of observed counts were converted into molecular weight via a mass calibration (**Supplementary Figure 3**), here with the 2G12 WT antibody monomer and dimer peaks. In the resulting molecular weight (MW) vs. counts histogram each species was identified as a resolved peak and counts were obtained from Gaussian fitting to these peaks. Molecular weights near the MP detection limit (ca. 40 kDa) were not quantified to ensure differentiation from background noise and were generally not treated as quantitative with respect to counts for larger species (**Supplementary Figure 12**). The measured counts were then converted into molar concentrations following the protocol described in equations **1-5** (for IgG and Fc*γ*RIa) and **equations 7-13** (for IgG and FcRn). For continuous flow experiments, the counts were diffusion-corrected by normalising to the MW-dependent factor of the diffusion coefficient (MW^−1/3^). For gasket experiments, the contribution from diffusion was negligible, as shown in **Supplementary Figure 4 & 5.**

#### *K*_d_ *screening experiments* **(Figure 2a-d and Figure 3a-c)**

*K*_d_ values are calculated by inserting the molar concentrations obtained from the previous section into **equation 6** for IgG-Fc*γ*RIa and **equations 14 –18** for IgG-FcRn.

#### *Kinetic experiments* **(Figure 2e & 2f, Supplementary Figure 20 & 21)**

Monitoring association or dissociation of the biomolecular complex allowed us to extract on-and off-rates. Additionally, we could extract *K*_d_ values after reaching equilibrium (plateau regions in the kinetic plots). Kinetic data sets were always composed of a series of individual MP experiments acquired at different time points after dilution from μM to pM-nM (continuous flow or gasket experiments). In both cases, we can extract the time-dependent ratio of free vs. bound species from each measurement at time *t* (e.g. **Figure 2e & 2f**). *Association experiment:* We begin by defining the differential equation describing the time-dependence for association (**equation 22-28**), based on the stoichiometric information from our MP experiments. In the association experiment, the starting concentrations of each species at *t* = 0 are known (**equations 19-21)**. We also know that [*IgG*_*bound*_]_0_ = 0 and [*IgG*_*bound*_] = *x*_*t*_, i.e. *x*_*t*_ is the amount of generated complex since mixing. The ordinary differential equation (ODE), **equation 28**, is numerically solved in Mathematica (Mathematica, Version 12.0) with the built-in function “Dsolve” to obtain x_t_ as a function of *k*_*on*_ and *k*_*off*_. The result for *x*_*t*_ is fit to the experimentally obtained ratio of the counts of IgG_bound_ and the total counts of IgG as described in **equation 29**. For fitting we used Mathematica’s built-in function “NonlinearModelFit” with the method “NMinimize” to optimize *k*_*on*_ and *k_off_*. From this we obtain *K_d_*, *k*_*on*_ and *k*_*off*_ values (**Supplementary Figure 20 & 21)**. *Dissociation experiment*: First, we calculate the *K*_*d*_ from the plateau region of the plot (**equations 1-6**). Knowing the *K*_*d*_ and the total concentrations of IgG and FcγIa before dilution (ca. μM), we can calculate the equilibrium concentrations of each species before dilution by solving **equation 32** and inserting into **equations 30 & 31**. At *t* = 0 after dilution, this distribution is unchanged, and the concentrations have to be divided by the dilution factor **(equations 33-35)**. This yields 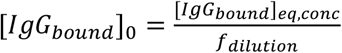 and [*IgG*_*bound*_] = [*IgG*_*bound*_]_0_ - *x*_*t*_, i.e. *x*_*t*_ is the amount of dissociated complex since diluting from the stock concentration to pM/nM concentration. Proceeding as described for the association experiment and using **equations 36-42** we obtain *K_d_*, *k*_*on*_ and *k*_*off*_ values (**Figure 2e & 2f**). In the future, we could combine the information from association and dissociation data and perform the fitting iteratively/simultaneously, from which *k*_off_, *k*_on_ and *K*_d_ can be obtained without waiting for equilibriation. This approach will extend our applicability to slow reactions by circumventing equilibration times of hours or even days (**Supplementary Figure 19**) and with this also reduce sample loss due to non-specific adsorption to sample tubes over time (**Supplementary Figure 22-24**).

### Protein Passivation of Sample Tubes

Bovine serum albumin (BSA, Sigma-Aldrich) was prepared as a 1 *μ*M solution in phosphate buffered saline (PBS) (Gibco® DPBS). 1 ml of this solution was added to each sample vial (1.5 ml Safe-Lock Tubes, Eppendorf) and rotated for ca. 2 hrs at room temperature (VWR rotator, VWR). The solution was completely removed, washed 3 times with PBS (1 ml each), filled with PBS and rotated for 30 min, then washed 3 times with PBS (1 ml each). All liquid was removed and replaced by either PBS buffer or 5 nM trastuzumab (Herceptin®). The trastuzumab solution was directly diluted in the passivated tube by mixing 2 *μ*l of trastuzumab with 398 *μ*l PBS. The control sample was prepared in the same way, but using an untreated sample vial.

Casein from bovine milk (technical grade, Sigma-Aldrich) was dissolved as 1 mg/ml in phosphate buffered saline (PBS) (Gibco® DPBS) at 42 C overnight. 1 ml of the solution was added to each sample vial (1.5 ml Safe-Lock Tubes, Eppendorf) and rotated for ca. 2 hrs at room temperature. The solution was completely removed, washed 3 times with PBS (1 ml each), filled with PBS and rotated for 30 min, then washed 3 times with PBS (1 ml each). All liquid was removed and replaced by either PBS buffer or 5 nM trastuzumab (Herceptin®). The trastuzumab solution was directly diluted in the passivated tube by mixing 2 *μ*l of trastuzumab with 398 *μ*l PBS. The control sample was prepared in the same way, but using an untreated sample vial.

### Non-specific Protein Adsorption to Sample Tubes

*m*PEG-silane coated glass vials (1.75 ml, Samco Trident Vial Tall with PP Screw Cap) were sonicated (PS-40A Ultrasonic Cleaner, Cole Palmer) for 10 min in 2% aqueous Hellmanex solution (Hellmanex III, HellmaAnalytics), washed with deionized water, then sonicated for 10 min in deionized water. The sonication step was repeated in ethanol. In a next step the glass vials were thoroughly dried under a stream of dry nitrogen and then placed in an O_2_ plasma (Zepto System, Diener Plasma Surface Technology) for 8 min. After plasma cleaning the vials were filled with a 5 mg/ml mPEG-silane solution in 1% acetic acid in ethanol. mPEG-silane was purchased from Laysan Bio as “MPEG-SIL-2000”. The vials were then placed in an oven (Vacucenter, SalvisLab) at 70 °C and the reaction solution mixed after 30, 60 and 90 min. After 2 hrs, the vials were emptied, washed with ethanol and deionized water and blow-dried under a stream of dry nitrogen. *Plasma cleaned glass vials*: The protocol for “*m*PEG-silane coated glass vials” was stopped after the plasma cleaning step. *Tween treated glass vials* (1.75 ml, Samco Trident Vial Tall with PP Screw Cap) were incubated for 4 hours with 2 % aqueous Tween solution (Tween20, Sigma-Aldrich), then rinsed with deion. water and blow-dried under a stream of dry nitrogen. For *untreated glass vials* we used 1.75 ml, Samco Trident Vial Tall with PP Screw Cap. Forprotein LoBind tubes we used Protein LoBind Tubes (1.5 ml, Eppendorf). Safe-Lock Tubes (1.5 ml, Eppendorf) were used as Eppendorf tubes. *Tween treated Protein LoBind* tubes (Protein LoBind Tubes (1.5 ml, Eppendorf)) were incubated for 4 hours with 2% aqueous Tween solution (Tween20, Sigma-Aldrich), then rinsed with deionized water and blow-dried under a stream of dry nitrogen. *PCR tubes* were Fisherbrand 0.2 ml PCR tubes.

In the last step, all tubes were washed with phosphate buffered saline (PBS) (Gibco® DPBS), dried and then incubated with 5 nM trastuzumab (Herceptin®) in PBS for 20 hrs at room temperature. A freshly prepared 5 nM IgG solution in LoBind tubes was used as a reference. The acquisition time of MP experiments was 90 s. Absolute count values of the IgG peak were then used to compare the performance of different sample tubes, i.e. counts were used as an indicator for absolute concentration.

### Surface Plasmon Resonance

Surface plasmon resonance experiments were carried out using a BIAcore T200 instrument (GE Healthcare). All experiments were performed in 10 mM HEPES, pH 7.4, 150 mM NaCl, 0.005% Tween 20 at 25 °C. Trastuzumab and deglycosylated trastuzumab mAb were immobilized on a CM5 chip (GE Healthcare) by amine coupling. Concentration series of human FcγRIa (A: 0.007-27 nM, B: 0.027-110 nM) were flowed over IgG (trastuzumab) (A) or deglycosylated IgG (trastuzumab) (B) bound surface at 30 μl/min for 120 s followed by buffer for 900 s. After each run, the biosensor chip was regenerated using 10 mM glycine, pH 2.5, which breaks the antibody-FcγR interaction. The specific binding response to antibody was obtained by subtracting the response given by analytes to an uncoupled surface and a blank run of buffer only. The kinetic sensorgrams were fitted to a global 1:1 interaction model to allow calculation of *k*_on_, *k*_off_, and *K*_d_ using BIAevaluation software 2.0.3 (GE Healthcare).

### Native Mass Spectrometry

Proteins were buffer exchanged into 1 M aqueous ammonium acetate (Sigma-Aldrich) using P6 Biospin columns (Bio-Rad) for the first two exchanges, followed by two exchanges using Amicon Ultra centrifugal filters (0.5 ml, 30 kDa MWCO) and in a last buffer exchange to 200 mM aqueous ammonium acetate using Amicon Ultra centrifugal filters. All protein solutions were analyzed at concentrations between 5 and 10 μM. Experiments were carried out on a prototype Thermo Scientific Q Exactive Hybrid Quadrupole Orbitrap Mass Spectrometer. Data were acquired using Xcalibur 3.0 software (Thermo Fisher Scientific) and raw spectra were deconvoluted to zero-charge spectra using UniDec^[36]^.

