## Supplementary Information for "Quantifying protein-protein interactions by molecular counting with mass photometry"

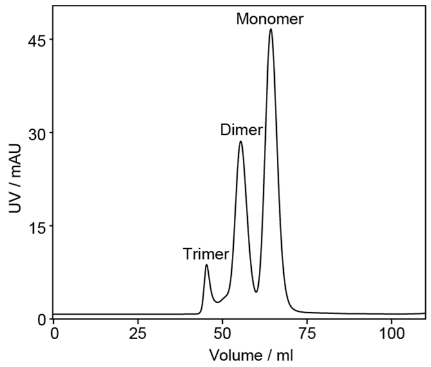

**Supplementary Figure 1: Size exclusion chromatography of 2G12.** UV profile of the monoclonal antibody 2G12 monomer, dimer and trimer complexes.

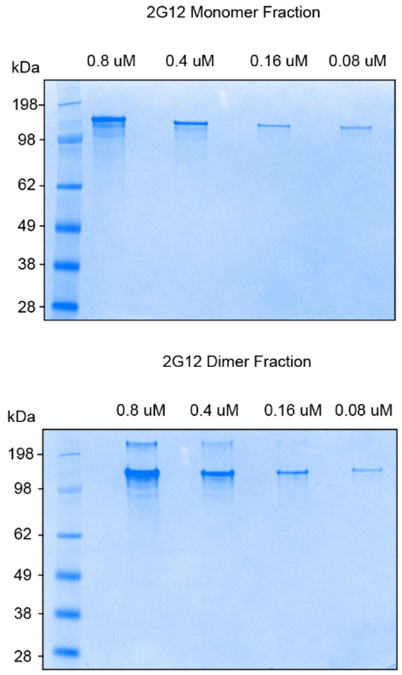

**Supplementary Figure 2: SDS-PAGE of SEC purified 2G12.** Monomers (top) and dimers (bottom). Concentrations from 0.8μM to 0.08 μM were imaged for each monomer/dimer SEC fraction.

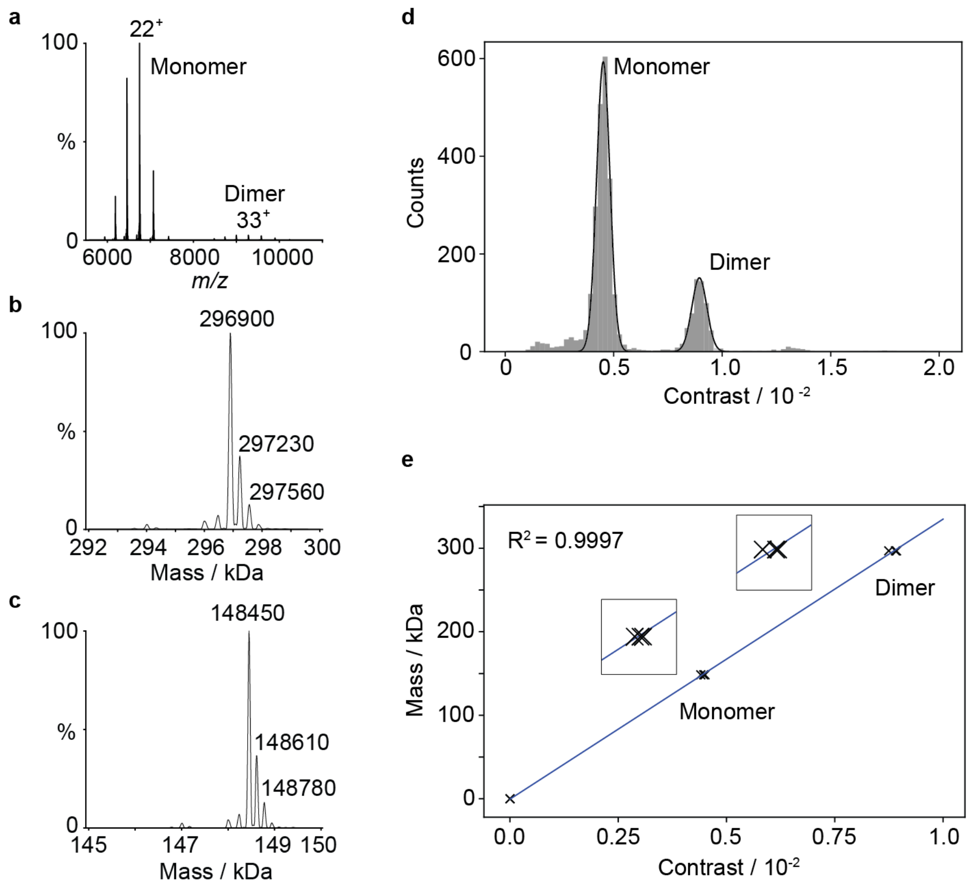

**Supplementary Figure 3: Native mass spectrometry of 2G12 and contrast calibration.** (**a**) Native mass spectrometry of 2G12 (top spectrum). (**b,c**) Zoom of the zero-charge state deconvoluted spectra, showing monomer mass range and dimer mass range. (**d**) Corresponding mass distribution obtained by MP. (**e**) MP mass calibration and reproducibility.

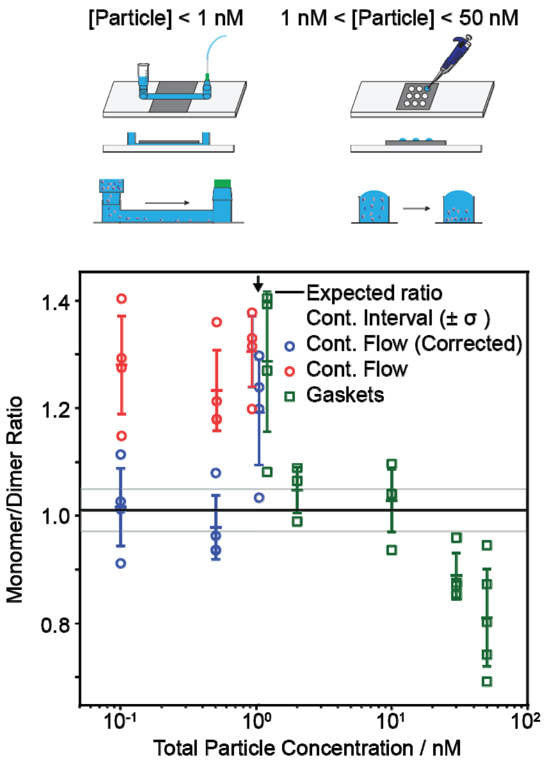

**Supplementary Figure 4: Dynamic range estimation, sample carriers and diffusion correction.** Experimental setup for continuous flow chambers (**top, left**) and silicone gaskets (**top, right**). MP ratio measurements of a 1:1 mixture of 2G12 monomer and dimer at different dilutions (0.1, 0.5, 1, 1, 2, 10, 30, 50 nM). The three results for 1 nM solutions were separated horizontally (**arrow**) to improve readability. Diffusion-corrected continuous flow ratios (blue) where obtained by normalizing monomer and dimer counts to their corresponding molecular weight dependent factor in the diffusion coefficient ($MW^{-1/3}$). The expected ratio (1.04 ± 0.05, black) was determined from UV-VIS measurements of concentrated ($\mu$M) stock solution of the pure monomer and dimer as well as weighing monomer and dimer stock solutions on a microbalance. These experiments were conducted to probe the dynamic range, suggesting that we can accurately measure ratios for samples with particle concentrations below 50 nM and extending the dynamic range to sub-nM concentrations when going from gaskets to a continuous-flow injection system. In the case of the continuous flow system we observed an influence of diffusion on our monomer:dimer ratios, which is expected to be molecular weight dependent ($MW^{-1/3}$). No influence was observed when conducting the experiments with gaskets when time between sample introduction and data recording was minimized (delay time <5 s) (**Supplementary Figure 5**).

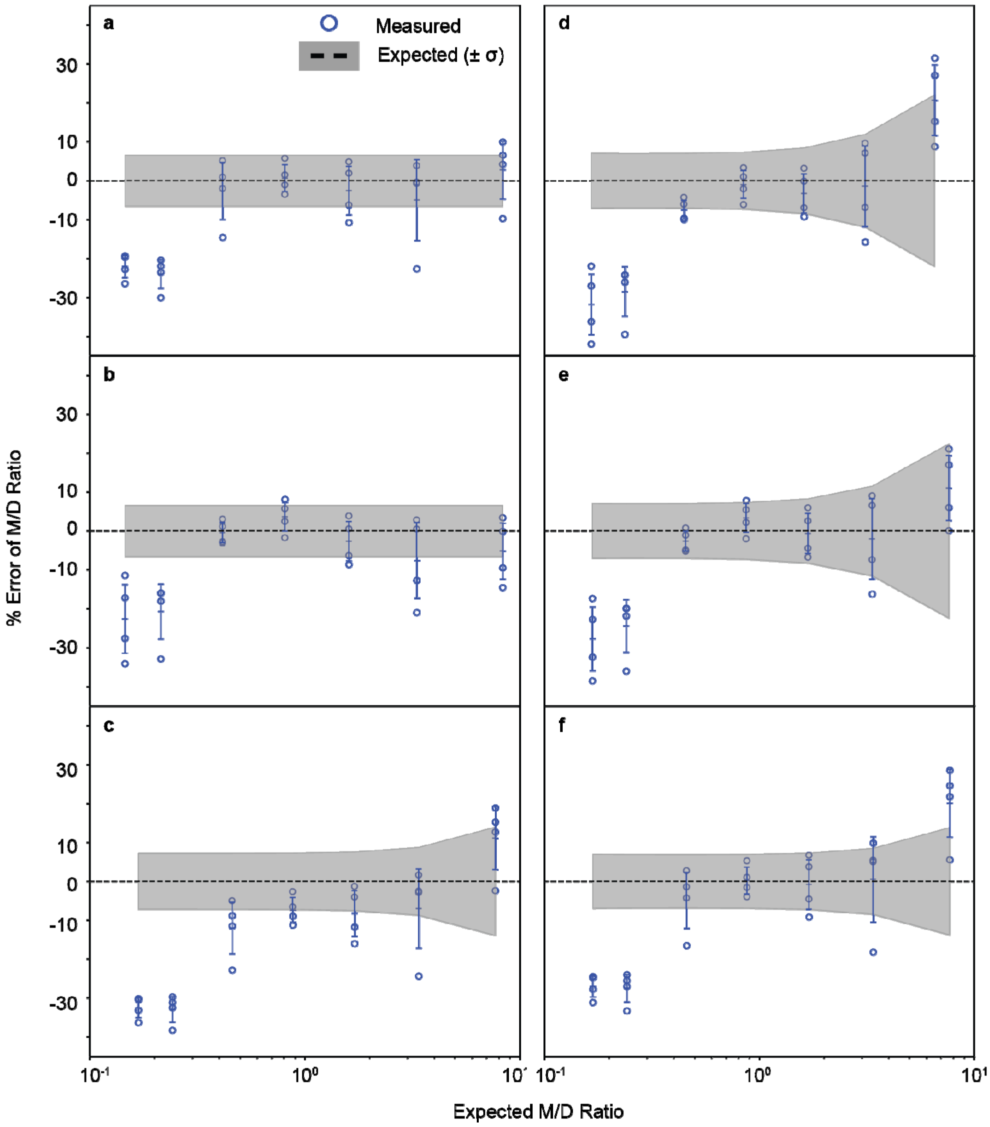

**Supplementary Figure 5: Data analysis approaches for 2G12 monomer/dimer ratios using sample gaskets.** We tested the influence of data acquisition duration, diffusion correction ($MW^{-1/3}$) and correcting for minor impurities. (**a**) No corrections, 90 s acquisition time. (**b**) No correction and 30 s acquisition time. The combination of accurate results, not needing to apply correction and having the highest time resolution were the main reasons to use method (**b**) in our work. (**c**) 90 s acquisition time and correction for minor impurities from control experiments of pure monomer, dimer and trimer. (**d**) 30 s acquisition time and correction for minor impurities from control experiments of pure monomer, dimer and trimer. (**e**) 90 s acquisition time and correction for minor impurities from control experiments of pure monomer, dimer and trimer and diffusion correction. (**f**) 30 s acquisition time and correction for minor impurities from control experiments of pure monomer, dimer and trimer and diffusion correction. We concluded that acquisition duration, diffusion correction and correction for minor impurities has only limited influence on the M:D ratio when the time between sample injection and acquisition start is kept below 5 seconds. From **Supplementary Figures 4 & 5**, we concluded that experimental procedures have to be carefully designed when working at sub-µM concentrations (e.g. when adding additional dilution steps) because of non-specific adsorption of protein to sample tube surfaces. Additionally, relative abundances obtained from continuous-flow injection should be diffusion-corrected. In the case of our injection procedure with gaskets, we show that sample diffusion, as well as minor small mass contaminants have minor effects on the 2G12 monomer:dimer ratio determined by molecular counting.

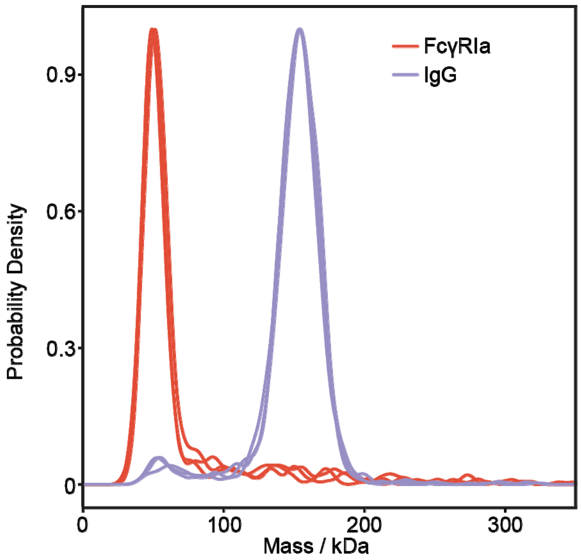

**Supplementary Figure 6: Technical replicates of purity screening of trastuzumab and FcγRIa.** Individual measurements of IgG (purple) were diluted from 7.4 µM to 5.2 nM and measured after 0.4, 3.8 and 7.6 min. Individual measurements of FcγIa (red) were diluted from 5 µM to 4.8 nM and measured after 0.4, 4.4 and 10.4 min. Small amounts of FcγRIa oligomers were visible, for IgG the data suggests high sample purity, with expected background at low molecular weight due to noise or fragments/residual impurities.

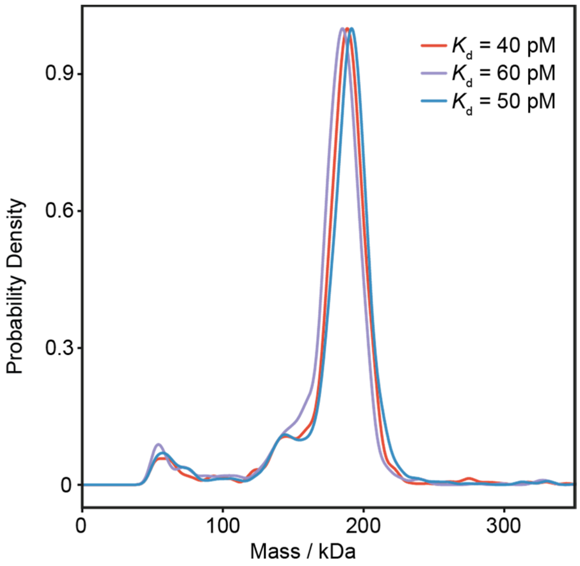

**Supplementary Figure 7: Technical replicates of IgG-FcγIa binding.** IgG-FcγIa were mixed 1:1 with final IgG concentration of 2.7 μM, followed by overnight incubation at room temperature. Samples were diluted to 4.4 nM IgG concentration and measured after 10, 14 and 18 min incubation time. Apparent $K_d$s in the low pM range were obtained. The small peak of free IgG at ca. 150 kDa could also originate from slighlty scewed 1:1 ratios due to uncertainity in the UV-VIS measurements.

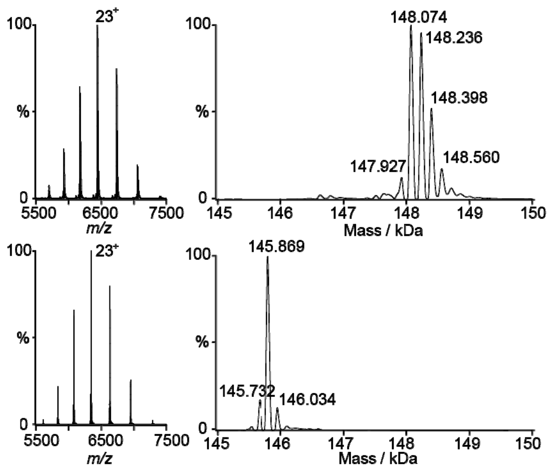

**Supplementary Figure 8: Native MS to confirm IgG deglycosylation.** Native MS of IgG (top, left) and corresponding zero-charge state deconvoluted spectrum (top, right). Corresponding deglycosylated IgG following treatment with Endoglycosidase S (native spectrum, bottom left and zero-charge state deconvoluted spectrum bottom, right).

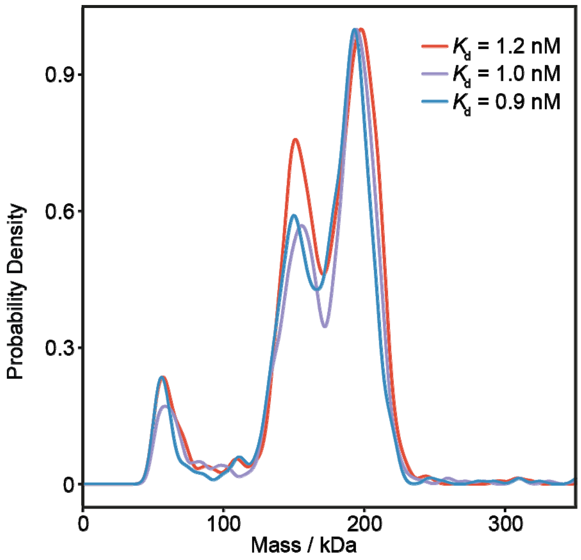

**Supplementary Figure 9: Technical replicates of $IgG_{deglycosylated}$-FcγIa binding.** $IgG_{deglycosylated}$ -FcγIa were mixed at a 1:1 ratio with final IgG concentration of 2.9 µM, followed by overnight incubation at room temperature. Samples were diluted to 4.6 nM IgG concentration and measured after 16, 13 and 18 min incubation time. Apparent $K_d$s in the low nM range were obtained.

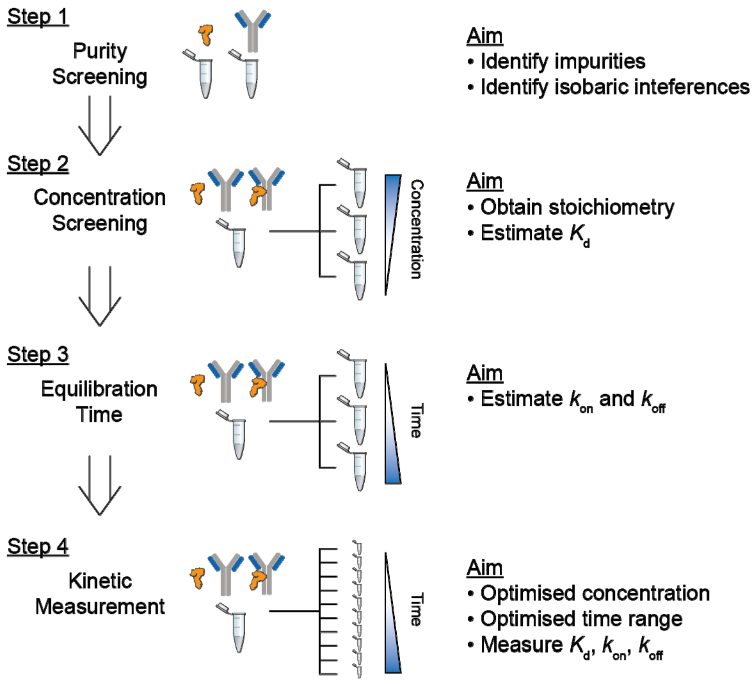

**Supplementary Figure 10: Screening method for quantifying binding affinities and kinetics with MP.** The ability to estimate binding affinities and kinetics in simple and fast screening experiments (concentration screening and equilibration time screening) offers a simple route to investigate a large number of candidates within a short time period and is crucial to prevent misinterpreting data derived from a single-shot $K_d$ approach. This allows us to choose ideal experimental parameters, such as concentration range and equilibration time, for a time-resolved experiment with which we can accurately determine the on and off rates of the interaction (**Supplementary Figure 18**).

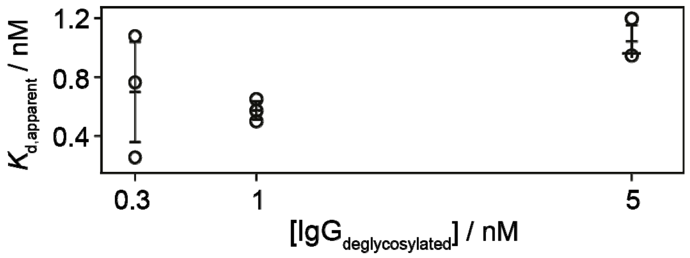

**Supplementary Figure 11: Concentration-dependent $K_d$ distribution for $IgG_{deglycosylated}$-FcγIa.** $IgG_{deglycosylated}$-FcγIa were mixed at a 1:1 ratio with a final IgG concentration of 2.9 μM, followed by overnight incubation at room temperature. Samples were diluted (0.3, 1.3 or 4.6 nM IgG concentration) and measured for 10 - 34 min incubation time. The time dependence of the apparent $K_d$ is best revealed for the 0.3 nM mixture (10 min at 0.3 nM: $K_d$= 0.25 nM, 22 min: $K_d$= 0.75 nM , 34 min: $K_d$= 1.1 nM, all at 0.3 nM).

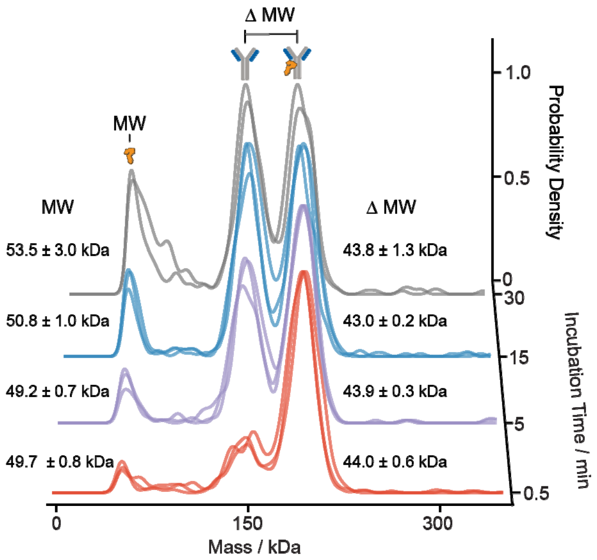

**Supplementary Figure 12: Equilibration time screening and assignment of FcγRIa molecular mass.** FcγRIa mass measured from the FcγRIa peak (left, around 50 kDa). FcγRIa mass measured from the mass difference between bound IgG (FcγRIa + IgG) and unbound IgG (right, 150- 200 kDa). We expect the molecular weight of FcγRIa (ca. 43-44 kDa) from the mass difference method to be more accurate than the direct read-out (49-54 kDa) due to its vicinity to the detection limit of our instrument. $IgG_{deglycosylated}$ -FcγIa were mixed at a 1:1 ratio with final IgG concentration of 2.9 μM, followed by overnight incubation at room temperature. Samples were diluted to 1.3 nM IgG concentration and measured after incubation times ranging from 0.4 - 30 min.

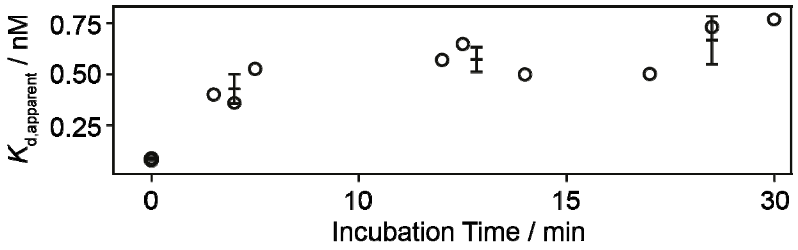

**Supplementary Figure 13: Equilibration time screening of $K_d$ for $IgG_{deglycosylated}$-FcγIa.** For FcγRIa binding to deglycosylated IgG, shifts in peak intensities were observed during the concentration screening (**Supplementary Figure 10**), indicating that we should be able to observe the time dependence of these interactions. Measurements revealed that equilibrium was reached after 10 minutes, yielding $K_d$ = 0.6 ± 0.1 nM (Figure 2d, Supplementary Figure 12). Repeating the same experiments (concentration and equilibration time screening) with FcγRIa-IgG highlighted the importance of this screening procedure (**Supplementary Figures 14-16**). For the here noted $IgG_{deglycosylated}$-FcγIa experiment we mixed the two compounds 1:1 with a final IgG concentration of 2.9 µM, followed by overnight incubation at room temperature. Samples were diluted to 1.3 nM and measured after 0.4-30 min incubation time (see **Supplementary Figure 12** for corresponding KDE-plots). Starting from apparent $K_d$ values of 0.09 nM after 0.4 min incubation time we can observe a gradual increase over time, until we start to reach a plateau-region after ca. 10 min, suggesting a $K_d$ = 0.6 ± 0.1 nM.

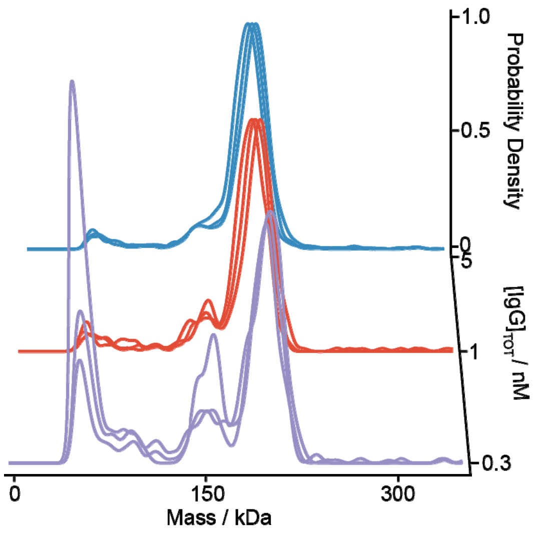

**Supplementary Figure 14: Concentration screening of IgG-FcγIa complexes.** IgG - FcγIa was mixed at a 1:1 ratio with final IgG concentration of 2.7 μM, followed by overnight incubation at room temperature. Samples were diluted (0.26, 1.3 and 4.4 nM) and measured after 9.6 - 18 min incubation time. With increasing dilution and increasing incubation time, we found a minimal increase in the unbound IgG peak intensity (apparent $K_d$ = 80 ± 10 pM at 4.4 nM, 25 ± 6 pM at 1.5 nM and 18 ± 7 at 260 pM). From this, a $K_d$ in the sub-nM range and an equilibration time >20 min can be estimated.

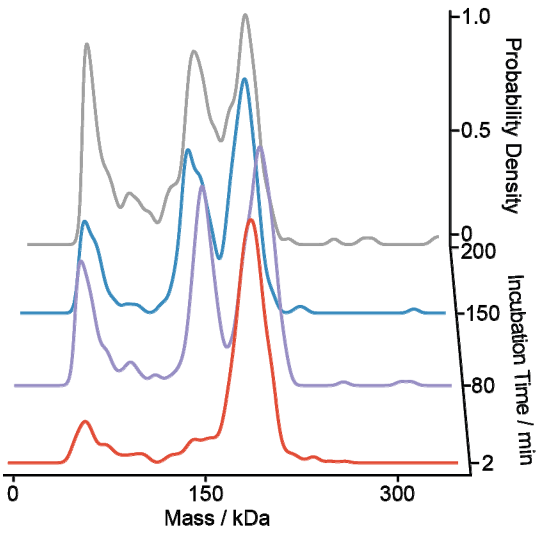

**Supplementary Figure 15: Equilibration time screening for IgG-FcγIa.** IgG -FcγIa were mixed at a 1:1 ratio with final IgG concentration of 2.7 µM, followed by overnight incubation at room temperature. Samples were diluted (0.28 and 0.21 nM) and measured after 2-200 min. Starting with an apparent $K_d$ value of 200 fM after 2 min, $K_d$ values are reaching a plateau-region after > 80 min incubation time, with $K_d$ values between 60-100 pM.

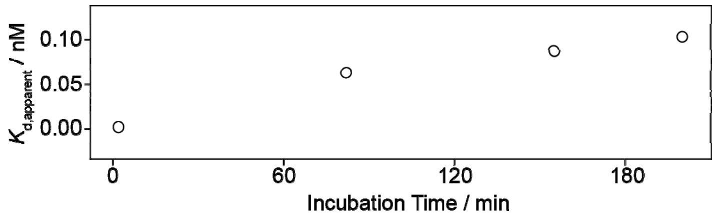

**Supplementary Figure 16: $K_d$ values from equilibration time screening for IgG-FcγIa.** Calculated from data in **Supplementary Figure 14**.

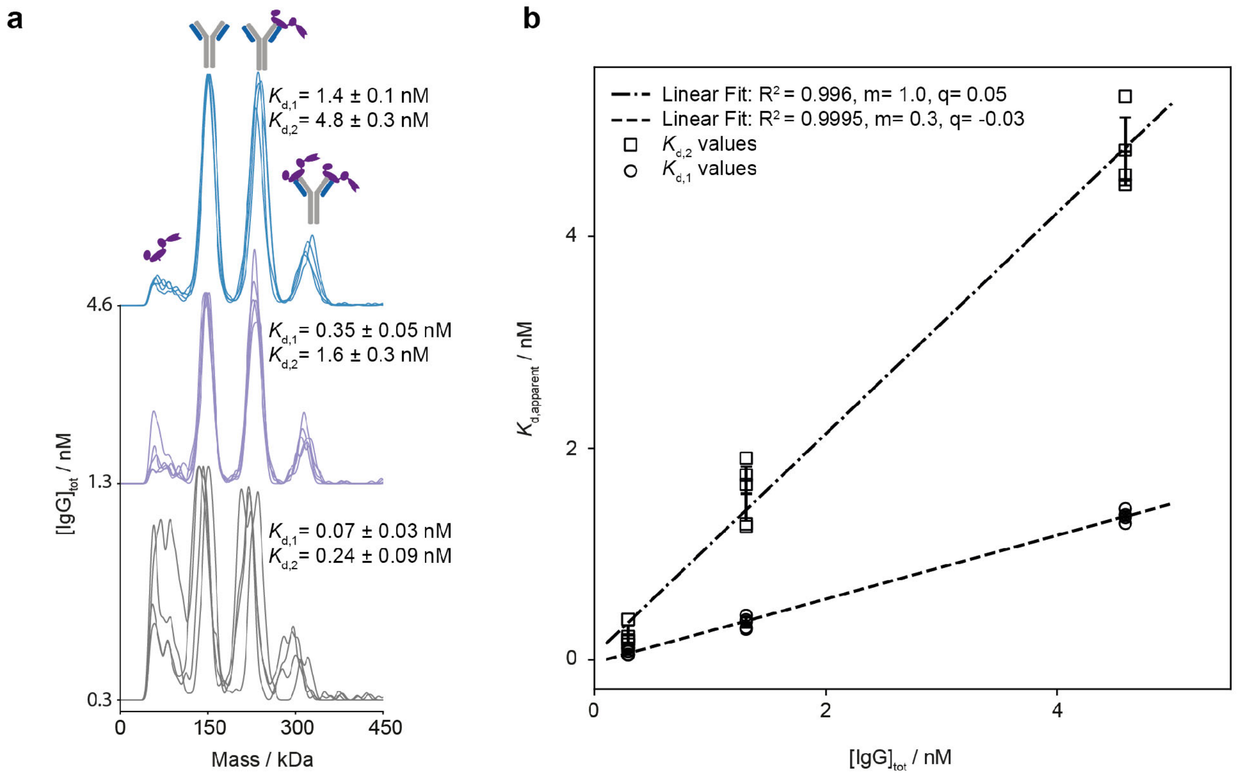

**Supplementary Figure 17: Screening the trastuzumab-HER2 interaction.** (**a**) 1.8 µM trastuzumab and 1.8 µM HER2 mixtures were equilibrated at room temperature overnight and diluted to desired pM-nM concentrations (0.3 nM, 1.3 nM and 4.6 nM). No significant differences in relative abundances of bound/unbound complexes were observed at various incubation times (2.5 – 100 min) for all concentrations. (**b**) Apparent $K_d$ values showed a strong linear dependence on concentration, indicating non-equilibrium conditions and/or $K_d$ values, which are exceeding our current dynamic concentration range/sensitivity of MP (i.e. sub-pM $K_d$). No valid $K_d$ value could be determined but the data suggests very strong binding affinities of trastuzumab to HER2 (approximately < pM) and/or very slow off rates (> hrs). This example highlights the importance of concentration and equilibration time screening for accurate $K_d$ measurements.

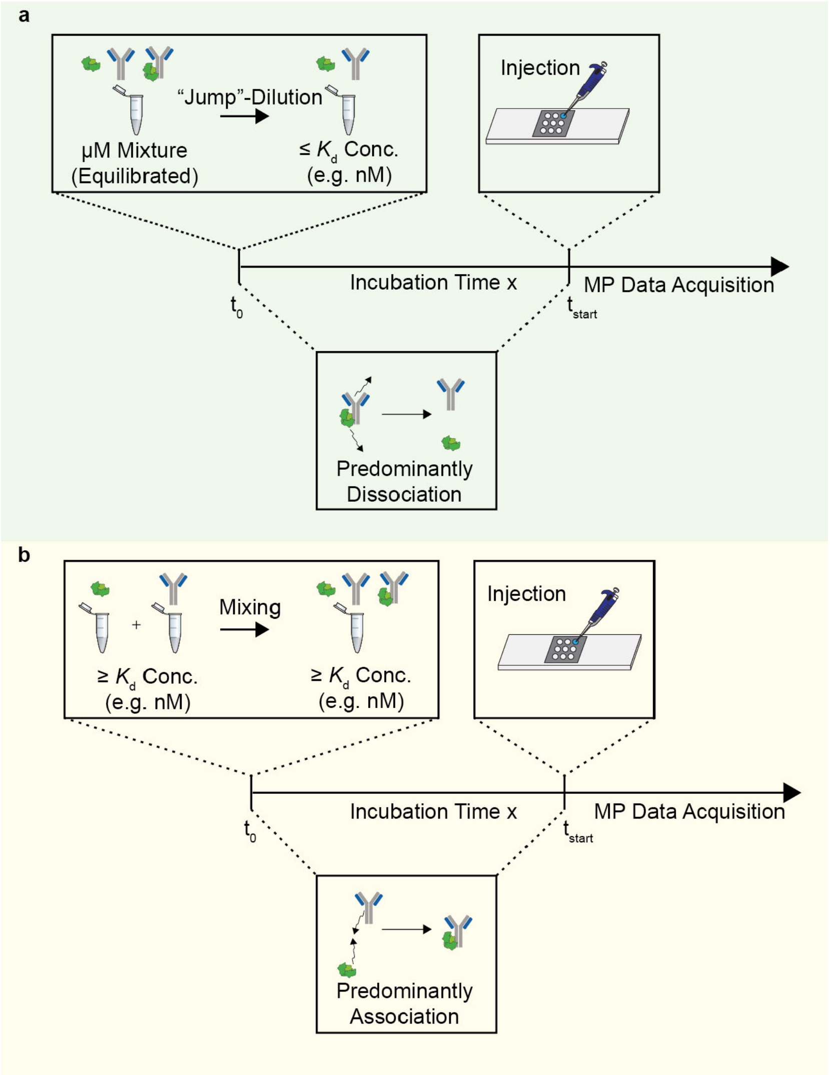

**Supplementary Figure 18: Principle of dissociation and association measurements.** A jump-dilution strategy for measuring $K_d$ and kinetics. Dilution to sub-$K_d$ concentrations results in dissociation of protein complexes over time (from $t_0$). After incubation time $t_{start}$ we quantify the bound to unbound ratio with a MP measurement, typically lasting 30 s (for gaskets). Mixing at concentrations above the expected $K_d$, in association experiments permits the quantification of complexes that form from $t_0$. Importantly, non-specific protein adsorption is a factor in both methods, but more noticeable for association measurements.

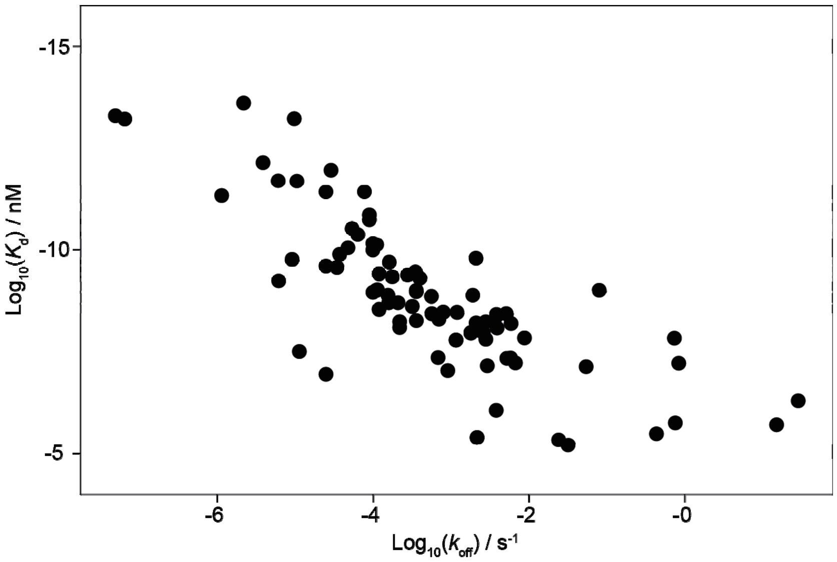

**Supplementary Figure 19: Correlation of published $K_d$ vs $k_{off}$ values.** Various biomolecular interactions measured by orthogonal techniques (e.g. SPR, BLI). Data is available in **Supplementary Table 1**. These data show a general correlation between stronger binding affinities corresponding to slower $k_{off}$ rates.

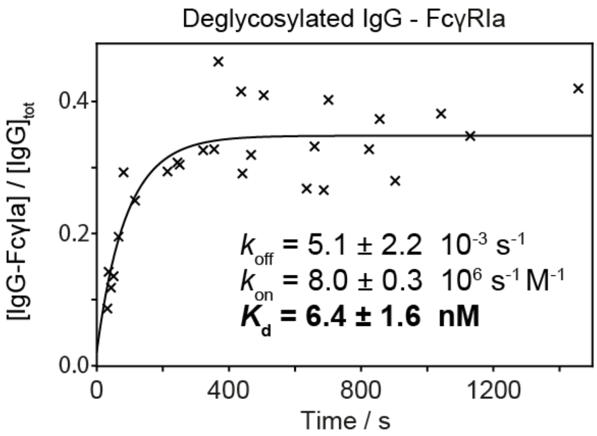

**Supplementary Figure 20: Association measurements ($k_{on}$) of $IgG_{deglycosylated}$-FcγIa.** $IgG_{deglycosylated}$-FcγIa was mixed at a 1:1 ratio with final concentrations of 4.9 nM for FcγIa and 4.6 nM for $IgG_{deglycosylated}$. Individual measurements (x) were taken after different incubation times (0.4 – 15 min). The $k_{on}$, $k_{off}$ and $K_d$ values were obtained from a non-linear fit (black line) to the experimental data and were in good agreement with values obtained from the dissociation experiment.

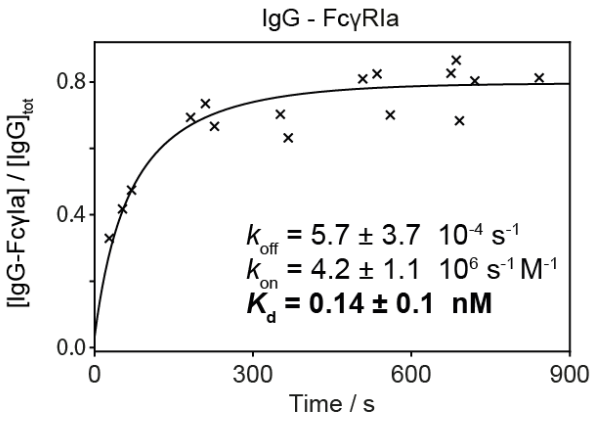

**Supplementary Figure 21: Association measurements ($k_{on}$) of IgG-FcγIa.** IgG-FcγIa were mixed at a 1:1 ratio with final concentrations of 3.0 nM for FcγIa and 3.1 nM for IgG. Individual measurements (x) were taken at different incubation times (0.5 – 44 min). The $k_{on}$, $k_{off}$ and $K_d$ values were obtained from a non-linear fit (black line) to the experimental data and were in good agreement with values obtained from the dissociation experiment.

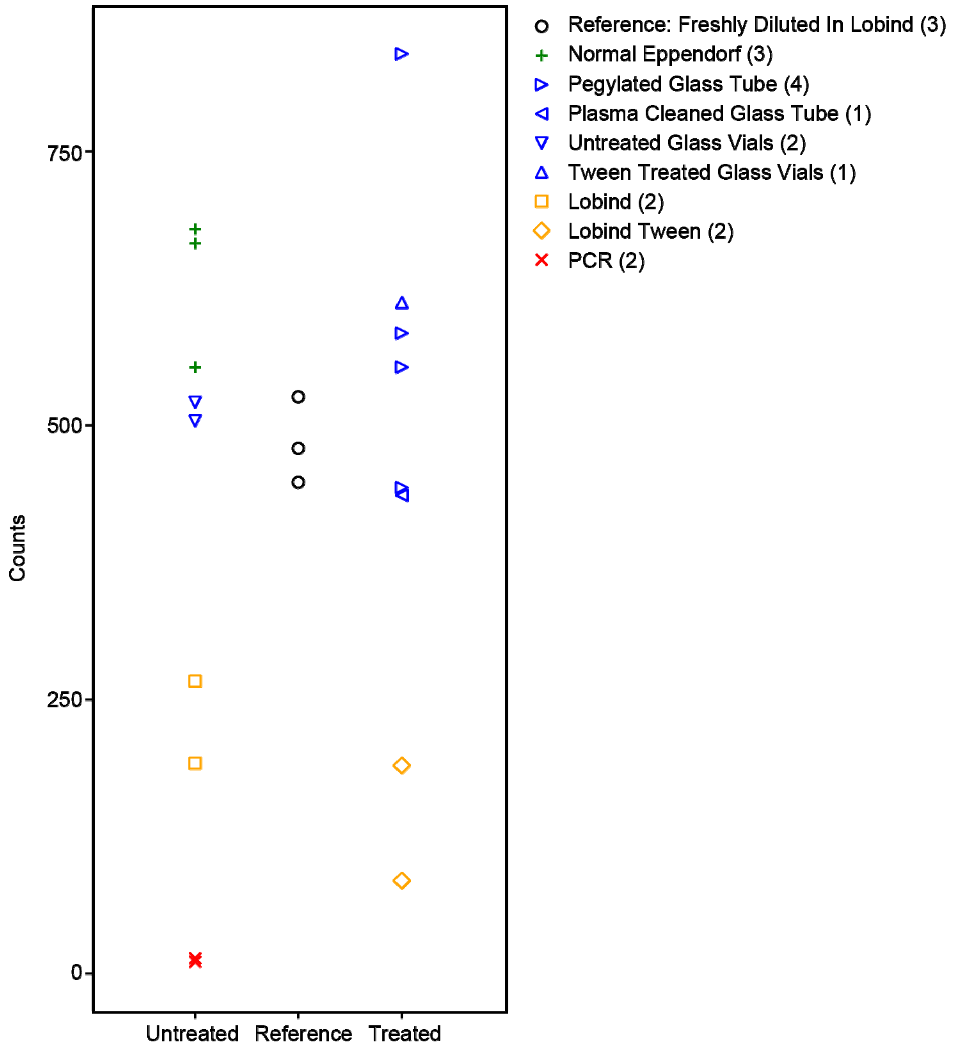

**Supplementary Figure 22: Non-specific IgG adsorption to samples tubes.** We freshly prepared 5 nM IgG in Eppendorf Lobind tubes to compare with the performance of other alternative materials, where IgG was stored for 20 hrs. Normal Eppendorf tubes were able to maintain the counts/concentration during this period. We saw good performance of glass vials, potentially due to their smaller surface-to-volume ratio. Eppendorf Lobind showed around 50% loss and PCR tubes >90% loss. We concluded from these results that normal Eppendorf tubes were most suitable for our experiments.

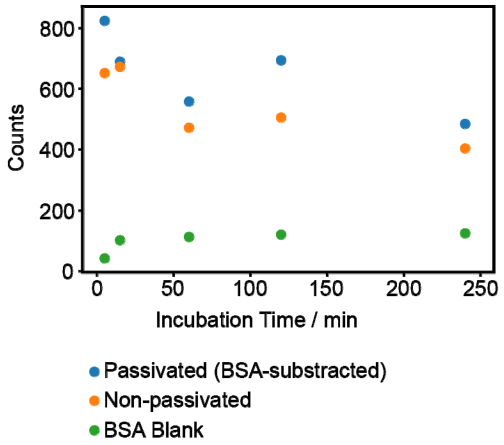

**Supplementary Figure 23: Protein passivation of sample tubes with BSA.** BSA showed no significant improvement to help maintaining concentration, i.e. counts, over time, potentially due to its high solubility.

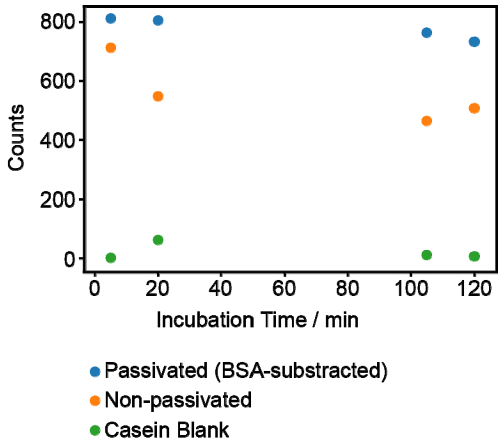

**Supplementary Figure 24: Protein passivation of sample tubes with casein.** Casein passivation helped maintain the concentration close to the initial level. Casein, seems to be ideal to passivate plastic surfaces, most likely because of its low solubility in water. Additionally, due to its low molecular weight (main species < 25 kDa), it does not cause interference in MP as it is below the detection limit of the current instrument.

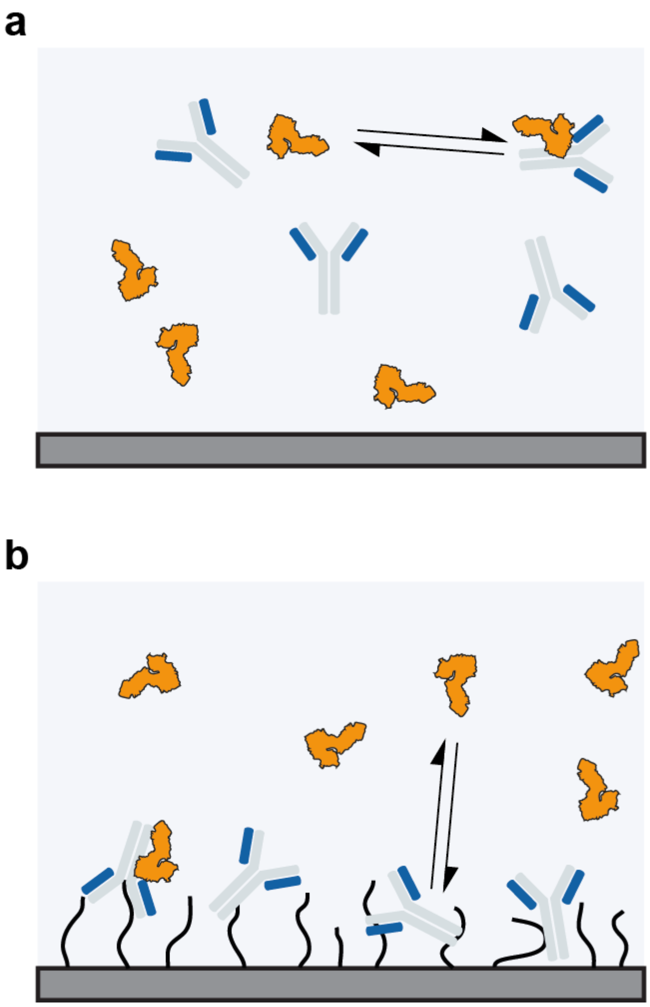

**Supplementary Figure 25: Schematic of interactions in MP and SPR.** (**a**) In-solution, label-free interactions occuring in MP. (**b**) Surface-immobilization (e.g. in a dextran matrix) and associated interactions in SPR. Differences in on-rates between SPR and MP (**Figure 2g & 2h** and **Supplementary Figure 20 & 21)** are attributed to mass transport, protein immobilization (i.e. orientation of IgG) and matrix effects [1]. Lower experimental on-rates, therefore, lead to greater calculated $K_d$ values by SPR.

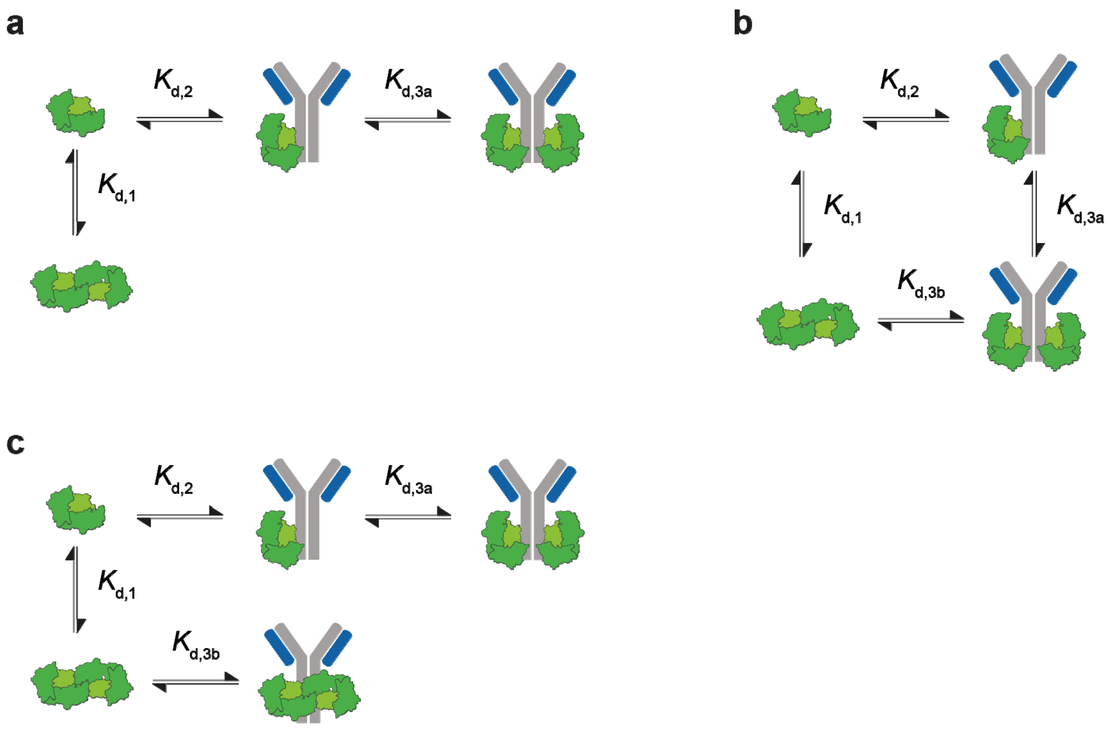

**Supplementary Figure 26: Proposed binding models for IgG-FcRn interactions.** Based on the existing literature we calculated binding affinities based on the free monomer binding model (a). The stoichiometries observed in our IgG-FcRn data would support also other models (**b, c**).

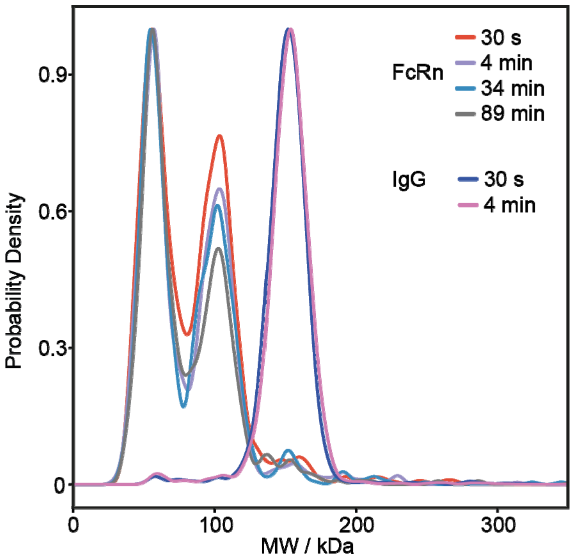

**Supplementary Figure 27: Technical replicates of IgG and FcRn pH = 5.** IgG (9 nM, purple) and FcRn (12 nM, red) measured at different time points. We can observe monomeric IgG at pH = 5 and FcRn present as monomer, dimer and in small quantities as trimer. The data suggests that FcRn reaches equilibrium within minutes (<30 min).

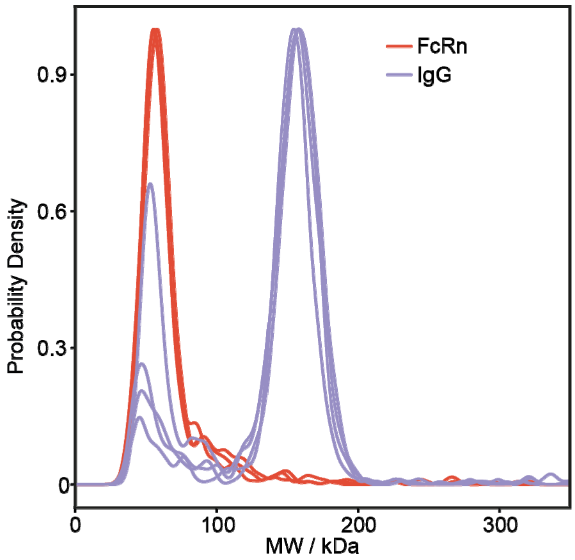

**Supplementary Figure 28: Technical replicates of IgG and FcRn pH = 5.5.** IgG (purple) was diluted from 3.6 µM to 2.5 nM and measured after different incubation times (0.4, 4.2, 10.9, 14.3 min). FcRn (red) was diluted from 7.1 µM to 4.5 nM and measured after 0.4, 4.2 and 15.7 min. Compared to pH = 5.0 we observe significantly less FcRn dimer.

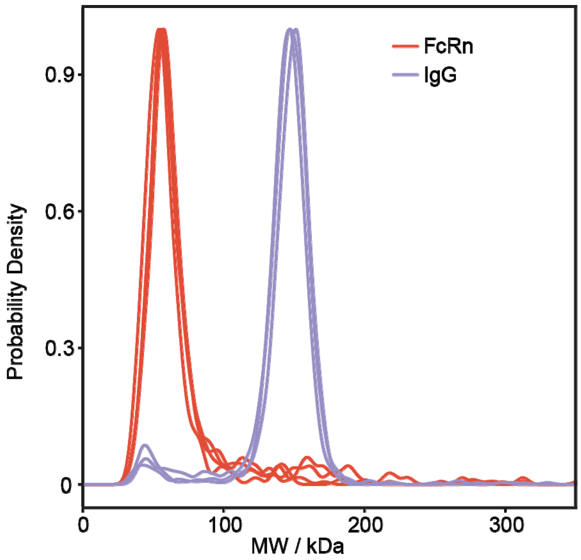

**Supplementary Figure 29: Technical replicates of IgG and FcRn pH = 6.0.** IgG (purple) was diluted from 2.0 μM to 4 nM and measured after different incubation times (0.7, 4.0, 7.0 min). FcRn (red) was diluted from 19.9 μM to 6.1 nM and measured after 0.4, 1, 3.8 and 12 min. Compared to pH = 5.0 we observe significantly less FcRn dimer.

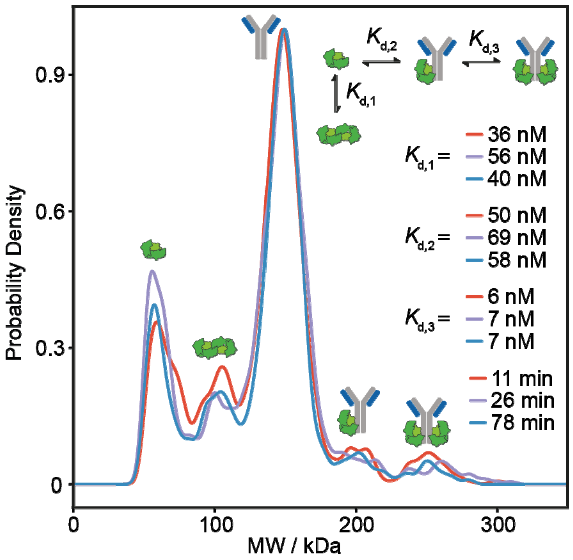

**Supplementary Figure 30: Time point and $K_d$ measurements of IgG-FcRn at pH = 5.** IgG concentrations were 3 nM IgG and 6 nM FcRn. Measurements were taken after 11 (red), 26 (purple) and 78 (blue) minutes equilibration time. The IgG-FcRn at pH 5.0 showed predominantly unbound IgG (88 ± 1 %), minor amounts of IgG bound to one (7 ± 1%) and two (5 ± 1%) FcRns.

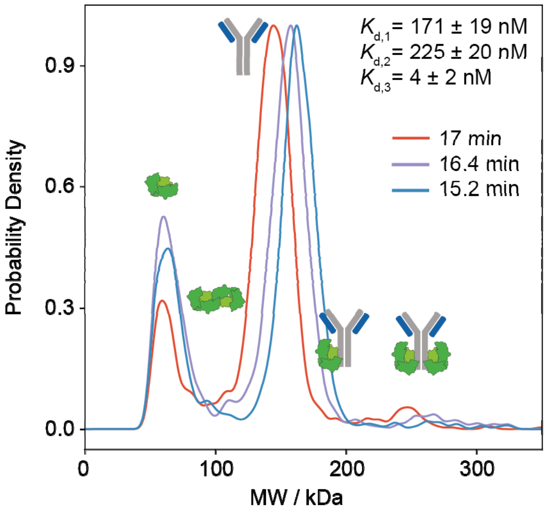

**Supplementary Figure 31: Time point and $K_d$ measurements of IgG-FcRn at pH = 5.5.** IgG concentrations were 3 nM IgG and 6 nM FcRn. Measurements were taken at 17 (red), 16 (purple) and 15 (blue) minutes.

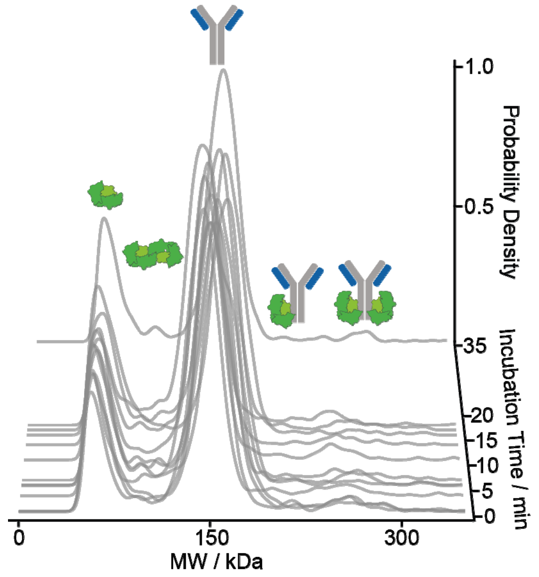

**Supplementary Figure 32: Time course measurements of IgG-FcRn binding at pH = 5.5.** Measurements were taken from 0.4 to 35 minutes at 3 nM IgG and 6 nM FcRn. Over the time course of 35 min no further dissociation of the bound species could be observed, suggesting that equilibrium is reached rapidly (<min).

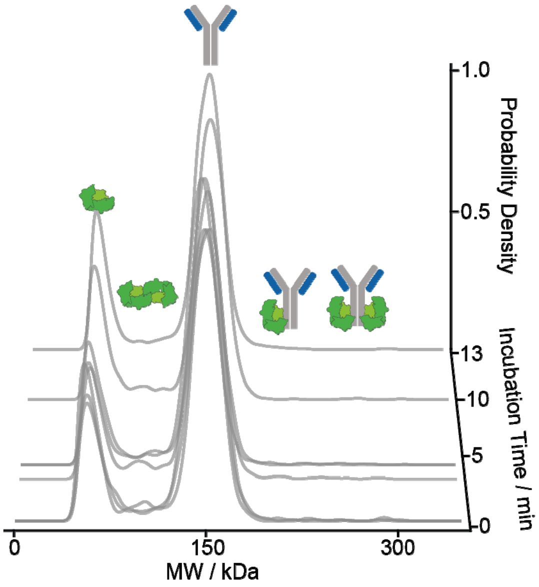

**Supplementary Figure 33: Time course measurements of IgG-FcRn binding at pH = 6.0.** Measurements were taken from 0.3 to 13 minutes at 3 nM IgG and 6 nM FcRn. Bound species were too low in abundance to be quantified. $K_d$ values are expected to be above 200 nM.

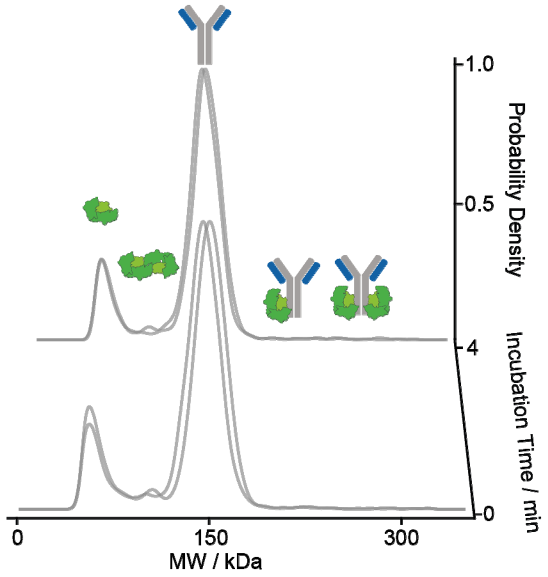

**Supplementary Figure 34: Time course measurements of IgG-FcRn binding at pH = 7.0.** Measurements were taken from 0.3 to 4 minutes at 3 nM IgG and 6 nM FcRn. Bound species were too low in abundance to be quantified. $K_d$ values are expected to be above 200 nM.

### 11. Supplementary Table 1: Raw data of published biomolecular binding affinities.

| #PDB | Affinity (M) | $k_{off}$ ($s^{-1}$) | Protein 1 | Protein 2 | Method | reference |
|---|---|---|---|---|---|---|
| 2FTL_E_I | 5.00E-14 | 5.00E-08 | Bovine trypsin | BPTI | IASP | [2] |
| 1TM1_E_I | 7.00E-13 | 3.90E-06 | Subtilisin BPN | Chymotrypsin inhibitor 2 | IASP | |
| 3W2D_A_HL | 5.79E-10 | 6.18E-06 | Staphylococcal enterotoxin B | 3E2 fab | SPR | |
| 1TM1_E_I | 2.00E-12 | 1.06E-05 | Subtilisin BPN | Chymotrypsin inhibitor 2 | SFFL | |
| 4HFK_A_BD | 2.69E-10 | 3.46E-05 | Tae4 | Tai4 | SPR | |
| 5C6T_HL_A | 1.30E-10 | 3.75E-05 | 1G2 fab | HCMV glycoprotein B | SPR | |
| 4U6H_AB_E | 9.00E-11 | 4.80E-05 | M12B9 fab | Vaccinia L1 | BI | |
| 3HFM_HL_Y | 3.00E-11 | 5.40E-05 | HyHEL-10 | HEW Lysozyme | | |
| 4CVW_A_C | 4.20E-11 | 6.40E-05 | Limit dextrinase | Limit dextrinase inhibitor | SPR | |
| 2SIC_E_I | 1.80E-11 | 9.00E-05 | Subtilisin BPN | Streptomyces subtilisin inhibitor | IASP | |
| 3HFM_HL_Y | 7.00E-11 | 1.00E-04 | HyHEL-10 | HEW Lysozyme | | |
| 2VIR_AB_C | 1.00E-09 | 1.10E-04 | IgG1 lambda fab | Flu virus hemagglutinin | SPR | |
| 3HFM_HL_Y | 7.50E-11 | 1.12E-04 | HyHEL-10 | HEW Lysozyme | | |
| 1BJ1_HL_VW | 2.90E-09 | 1.20E-04 | Fab-12 | VEGF | | |
| 1N8Z_AB_C | 1.31E-09 | 1.56E-04 | Herceptin | erbB-2 | SPR | |
| 4MNQ_ABC_DE | 2.00E-09 | 1.60E-04 | HLA-A2 plus telomerase peptide | ILA1 TCR | SPR | |
| 3BT1_A_U | 4.60E-10 | 1.77E-04 | Urokinase-type plasminogen activator | Urokinase plasminogen activator surface receptor | SPR | |
| 1GL0_E_I | 2.00E-09 | 2.10E-04 | Bovine alpha-chymotrypsin | PMP-D2v insect inhibitor | | |
| 4K71_A_BC | 8.00E-09 | 2.20E-04 | Human Serum Albumin | FcRn | | |
| 3NGB_HL_G | 5.76E-09 | 2.20E-04 | VRC01 fab | gp120 | SPR | |
| 2DSQ_I_G | 4.14E-10 | 2.78E-04 | IGF-I | IGFBP1 | SPR | |
| 4HSA_AB_C | 2.46E-09 | 3.20E-04 | Interleukin-17a | Interleukin-17 receptor A | SPR | |
| 1N8Z_AB_C | 3.50E-10 | 3.50E-04 | Herceptin | erbB-2 | SPR | |
| 2B42_A_B | 1.07E-09 | 3.60E-04 | TAXI-I | B. subtilis endoxylanase | SPR | |
| 1N8Z_AB_C | 5.00E-10 | 4.00E-04 | Herceptin | erbB-2 | SPR | |
| 1DAN_HL_UT | 3.70E-09 | 5.70E-04 | Factor VIIa | Tissue factor | | |
| 4JPK_HL_A | 4.36E-08 | 6.83E-04 | VRC01 fab | eOD-GT6 | SPR | |
| 2AJF_A_E | 1.62E-08 | 1.16E-03 | Human Angiotensin-converting enzyme 2 | SARS spike protein receptor binding domain | SPR | |
| 1CBW_FGH_I | 1.10E-08 | 1.80E-03 | Bovine alpha-chymotrypsin | BPTI | IASP | |
| 1YY9_CD_A | 1.31E-09 | 1.91E-03 | Cetuximab fab | Epidermal growth factor receptor | SPR | |
| 2I26_N_L | 9.50E-09 | 2.00E-03 | Type II IgNAR | HEW Lysozyme | SPR | |
| 1IAR_A_B | 1.62E-10 | 2.10E-03 | Interleukin-4 | Interleukin-4 receptor | SPR | |
| 1DAN_HL_UT | 6.16E-09 | 2.10E-03 | Factor VIIa | Tissue factor | | |
| 2VIS_AB_C | 4.00E-06 | 2.16E-03 | IgG1 lambda fab | Flu virus hemagglutinin | | |
| 1MHP_HL_A | 1.07E-08 | 2.60E-03 | AQC2 fab | Integrin alpha-1 | SPR | |
| 1WQJ_I_B | 5.78E-09 | 2.79E-03 | IGF-I | IGF-1R | SPR | |

| | | | | | | |
|---|---|---|---|---|---|---|
| 3BP8_A_C | 4.14E-09 | 3.85E-03 | Mlc transcription regulator | PTS glucose-specific enzyme EIICB | SPR | |
| 1NMB_N_LH | 4.55E-08 | 5.20E-03 | Subtype N9 neuraminidase | Antibody NC10 | SPR | |
| 5F4E_A_B | 5.90E-08 | 6.70E-03 | Sperm-egg fusion protein Izumo | Sperm-egg fusion protein Juno | BI | |
| 1JRH_LH_I | 1.44E-08 | 8.75E-03 | mAbs A6 | Interferon gamma receptor | SPR | |
| 4K71_A_BC | 6.20E-06 | 3.20E-02 | Human Serum Albumin | FcRn | | |
| 2ptc | 6.00E-14 | 6.60E-08 | Trypsinogen | BPTI | spectrophotometric assays | [3] |
| 1jiw | 4.60E-12 | 1.15E-06 | Alkaline metallo-proteinase | Proteinase inhibitor | spectrophotometric assays | |
| 1emv | 2.44E-14 | 2.20E-06 | Colicin E9 nuclease | Im9 immunity protein | stopped-flow fluorescence | |
| 2sni | 1.97E-12 | 6.10E-06 | Subtilisin | Chymotrypsin inhibitor 2 | spectrophotometric assays | |
| 1t6b | 1.74E-10 | 9.20E-06 | Anthrax protective antigen | Anthrax toxin receptor | SPR and FRET | |
| 1dfj | 5.90E-14 | 9.80E-06 | Ribonuclease A | Rnase inhibitor | RNAse assay | |
| 1ffw | 3.10E-08 | 1.14E-05 | Chemotaxis protein CheY | Chemotaxis protein CheA | SPR | |
| 1fsk | 2.50E-10 | 2.50E-05 | Fab - Birch pollen antigen Bet V1 | Birch pollen antigen Bet V1 | | |
| 1jmo | 1.12E-07 | 2.50E-05 | Heparin cofactor | Thrombin | spectrophotometric assays | |
| 1ppe | 3.73E-12 | 2.50E-05 | Trypsinogen | CMTI-1 squash inhibitor | spectrophotometric assays | |
| 1mah | 1.07E-12 | 2.90E-05 | Acetylcholinesterase | Fasciculin | spectrophotometric assays | |
| 1eer | 3.71E-12 | 7.80E-05 | Erythropoietin | EPO receptor | SPR | |
| 2sic | 1.38E-11 | 9.00E-05 | Subtilisin | Streptomyces subtilisin inhibitor | spectrophotometric assays | |
| 1jps | 1.02E-10 | 0.0001 | Fab D3H44 | Tissue factor | spectrophotometric assays | |
| 2i25 | 1.11E-09 | 0.0001 | Shark single domain antigen receptor | HEW lysozyme | SPR | |
| 2vir | 1.00E-09 | 0.00011 | Fab | Flu virus hemagglutinin | SPR | |
| 2i9b | 9.58E-10 | 0.000114 | uPAR surface receptor | Urokinase-type plasminogen activator | SPR | |
| 1jtg | 3.87E-10 | 0.00012 | β-lactamase inhibitor protein | β-lactamase TEM-1 | spectrophotometric assays | |
| 1gl1 | 2.03E-10 | 0.000162 | Chymotrypsin | PMP-C (LCMI II) | spectrophotometric assays | |
| 1jwh | 5.41E-09 | 0.00036 | Casein kinase II β chain | Casein kinase II α chain | SPR | |
| 2b42 | 9.97E-10 | 0.00036 | Xylanase | Xylanase inhibitor | SPR | |
| 2gox | 1.39E-09 | 0.000563 | Complement C3d fragment | Staphylococcus aureus Efb-C | SPR | |
| 1gxd | 5.00E-09 | 0.0007 | ProMMP2 type IV collagenase | Metalloproteinase inhibitor 2 | SPR | |
| 1kxq | 3.39E-09 | 0.0008 | Camel VHH - Pancreatic α-amylase | Pancreatic α-amylase | IAsys | |
| 1mlc | 9.10E-08 | 0.00091 | Fab44.1 | HEW lysozyme | SPR | |
| 2ajf | 1.63E-08 | 0.00116 | Angiotensin-converting enzyme 2 | SARS spike protein receptor binding domain | SPR | |
| 1e6j | 3.43E-09 | 0.0012 | Fab 13B5 | HIV-1 capsid protein p24 | SPR | |
| 1cbw | 1.06E-08 | 0.0018 | Chymotrypsin | BPTI | spectrophotometric assays | |
| 2vis | 4.00E-06 | 0.00216 | Fab | 1GIG LH | SPR | |
| 1kac | 1.56E-08 | 0.0028 | Adenovirus fiber knob protein | Adenovirus receptor | SPR | |
| 1p2c | 6.94E-08 | 0.00292 | FabF10.6.6 | HEW lysozyme | IAsys | |
| 1e6e | 8.57E-07 | 0.0038 | Adrenoxin reductase | Adrenoxin | SPR | |

| | | | | | |
|---|---|---|---|---|---|
| 3bp8 | 3.87E-09 | 0.00385 | Mlc transcription regulator | PTS glucose-specific enzyme EIICB | SPR |
| 2b4j | 8.21E-09 | 0.0039 | Integrase (HIV-1) | PC4 and SFRS1 interacting protein | fluorescence assay |
| 1vfb | 3.70E-09 | 0.00514 | Fv D1.3 | HEW lysozyme | stopped-flow fluorescence |
| 1kkl | 4.46E-08 | 0.0058 | HPr kinase C-ter domain | HPr | SPR |
| 1xu1 | 6.39E-09 | 0.00589 | TNF domain of APRIL | TACI CRD2 domain | SPR |
| 1klu | 4.60E-06 | 0.0239999 9 | MHC class 2 HLA-DR1 | Staphylococcus enterotoxin C3 | SPR |
| 1ktz | 7.30E-08 | 0.0540000 3 | TGF-β | TGF-β receptor | SPR |
| 2oza | 9.88E-10 | 0.08 | MAP kinase 14 | MAP kinase-activated protein kinase 2 | stopped-flow fluorescence |
| 1mq8 | 3.23E-06 | 0.4299995 5 | ICAM-1 domain 1-2 | Integrin α-L I domain | SPR |
| 2wpt | 1.46E-08 | 0.7300002 4 | Colicin E9 nuclease | Im2 immunity protein | stopped-flow fluorescence |
| 1e4k | 1.74E-06 | 0.7499995 5 | FC fragment of human IgG 1 | Human FCGR III | SPR |
| | 6.00E-08 | 0.83 | Colicin E9 nuclease | Im2 immunity protein | stopped-flow fluorescence |
| 1lfd | 1.94E-06 | 14.899990 8 | Ras.GNP | RalGDS Ras-interacting domain | stopped-flow fluorescence |
| | 5.00E-07 | 28.2 | Colicin E9 nuclease | Im8 immunity protein | stopped-flow fluorescence |

Index: EI (enzyme & inhibitor), ES (enzyme & substrate), ER (enzyme & receptor), OR (other & receptor), OX (other & miscellaneous), AB (antibody & antigen), SPR (surface plasmon resonance), IAsys (resonant mirror biosensor) , FRET (Förster resonance energy transfer microscopy), BI (biolayer interferometry), SFFL (stopped flow fluorescence).

### 12. Equations

Calculation of $K_d$ values from relative abundances measured with MP.
For the interactions of IgG - Fcγla and deglycosylated IgG- Fcγla we obtain the following equations:
Mass balance:

*(1)* $$[IgG]_{total} = [IgG]_{unbound} + [IgG]_{bound}$$

$[IgG]_{total}$ , $[IgG]_{unbound}$ and $[IgG]_{bound}$ are IgG molar concentrations

*Calculation of conversion factor:*

*(2)* $$f_{conversion} = \frac{[IgG]_{total}}{counts(IgG_{unbound})+counts(IgG_{bound})}$$

where $counts(IgG_{unbound})$ and $counts(IgG_{bound})$ are the counts obtained from Gaussian fits to the mass histograms, $f_{converesion}$ is the conversion factor which converts counts into molar concentrations

Conversion of counts to molar concentrations:

*(3)* $$[\mathrm{IgG}]_{\mathrm{bound}} = \mathrm{counts}\ (\mathrm{IgG}_{\mathrm{bound}}) * f_{conversion}$$

*(4)* $$[\mathrm{IgG}]_{\mathrm{unbound}} = \mathrm{counts}\ (\mathrm{IgG}_{\mathrm{unbound}}) * f_{conversion}$$

*(5)* $$[\mathrm{Fc\gamma Ia}]_{\mathrm{unbound}} = [\mathrm{Fc\gamma Ia}]_{\mathrm{total}} - [\mathrm{IgG}]_{\mathrm{bound}}$$

Due to variable noise contributions in the low molecular weight range (<80 kDa), we have to calculate the molar concentration of Fcγla via **equation 5**. The $K_d$ is then obtained from **equation 6**.

*(6)* $$K_\mathrm{d} = \frac{[\mathrm{Fc\gamma Ia}]_{\mathrm{unbound}} * [\mathrm{IgG}]_{\mathrm{unbound}}}{[\mathrm{IgG}]_{\mathrm{bound}}}$$

For the interactions of IgG-FcRn we obtain the following equation:

Mass balance:

*(7)* $$[IgG]_{total} = [IgG]_{unbound} + [IgG]_{single\ bound} + [IgG]_{double\ bound}$$

Calculation of conversion factor:

*(8)* $$f_{converesion} = \frac{[IgG]_{total}}{counts(IgG_{unbound})+counts(IgG_{single\ bound})+counts(IgG_{double\ bound})}$$

Conversion of counts to molar concentrations:

*(9)* $$[\mathrm{IgG}]_{\mathrm{bound}} = \mathrm{counts}\ (\mathrm{IgG}_{\mathrm{bound}}) * f_{conversion}$$

*(10)* $$[\mathrm{IgG}]_{\mathrm{single\ bound}} = \mathrm{counts}\ (\mathrm{IgG}_{\mathrm{single\ bound}}) * f_{conversion}$$

*(11)* $$[\mathrm{IgG}]_{\mathrm{double\ bound}} = \mathrm{counts}\ (\mathrm{IgG}_{\mathrm{double\ bound}}) * f_{conversion}$$

*(12)* $$[FcRn_{dimer}] = \mathrm{counts}\ (\mathrm{FcRn}_{\mathrm{dimer}}) * f_{conversion}$$

*(13)* $$[FcRn_{monomer}] = [FcRn]_{total} - 2 * [FcRn_{dimer}] - [\mathrm{IgG}]_{\mathrm{single\ bound}} - 2 * [\mathrm{IgG}]_{\mathrm{double\ bound}}$$

Calculation of $K_d$ values for interactions:

(14) $$K_1 = \frac{[FcRN_{monomer}]^2}{[FcRN_{dimer}]}$$

(15) $$K_2 = \frac{[FcRN_{monomer}]*[IgG]_{unbound}}{[\mathrm{IgG}]_{\text{single bound}}}$$

(16) $$K_{3a} = \frac{[FcRN_{monomer}]*[\mathrm{IgG}]_{\text{single bound}}}{[\mathrm{IgG}]_{\text{double bound}}}$$

(17) $$K_{3b} = \frac{[FcRN_{dimer}]*[IgG]_{unbound}}{[\mathrm{IgG}]_{\text{double bound}}}$$

(18) $$K_{3,app} = K_{3a} * K_{3b}$$

Kinetic experiments of IgG - FcγIa and deglycosylated IgG- FcγIa:

For association:

At t=0:

(19) $$[\mathrm{Fc\gamma Ia}] = [\mathrm{Fc\gamma Ia}]_0 = [\mathrm{Fc\gamma Ia}]_{tot}$$

(20) $$[IgG_{unbound}] = [IgG_{unbound}]_0 = [IgG]_{tot}$$

(21) $$[IgG_{bound}] = [IgG_{bound}]_0 = 0$$

At t>0:

(22) $$[IgG_{unbound}] = [IgG_{unbound}]_0 - x_t$$

(23) $$[\mathrm{Fc\gamma Ia}] = [\mathrm{Fc\gamma Ia}]_0 - x_t$$

(24) $$[IgG_{bound}] = [IgG_{bound}]_0 + x_t$$

(25) $$K_d = \frac{k_{off}}{k_{on}}$$

(26) $$\frac{d[IgG_{bound}]}{dt} = k_{on} * [\mathrm{Fc\gamma Ia}] * [IgG_{unbound}] - k_{off} * [IgG_{bound}]$$

(27) $$\frac{d[IgG_{unbound}]}{dt} = \frac{d[\mathrm{Fc\gamma Ia}]}{dt} = -\frac{d[IgG_{bound}]}{dt}$$

(28) $$\frac{dx}{dt} = k_{on}([\mathrm{Fc\gamma Ia}]_0 - x_t) * ([IgG_{unbound}]_0 - x_t) - k_{off} * ([IgG_{bound}]_0 + x_t)$$

(29) $$\frac{counts(IgG_{bound})}{counts(IgG_{tot})} = \frac{x_t}{[IgG]_{tot}}$$

For dissociation:
Calculation of the concentrations after equilibration of stock mixtures (ca. µM concentrations):

(30) $$[IgG_{unbound}]_{eq,conc} = [\text{Fc}\gamma\text{Ia}]_{eq,conc}$$

(31) $$[IgG_{unbound}]_{eq,conc} = [IgG]_{tot,conc} - [IgG_{bound}]_{eq,conc}$$

(32) $$K_d = \frac{([IgG]_{tot,conc} - [IgG_{bound}]_{eq,conc})^2}{[IgG_{bound}]_{eq,conc}}$$

Using the known dilution factor (from dilutiong the stock mixtures (ca. µM) to nM/pM concentrations):

At t=0:

(33) $$[\text{Fc}\gamma\text{Ia}] = [\text{Fc}\gamma\text{Ia}]_0 = \frac{[\text{Fc}\gamma\text{Ia}]_{eq,conc}}{f_{dilution}}$$

(34) $$[IgG_{unbound}] = [IgG_{unbound}]_0 = \frac{[IgG_{unbound}]_{eq,conc}}{f_{dilution}}$$

(35) $$[IgG_{bound}] = [IgG_{bound}]_0 = \frac{[IgG_{bound}]_{eq,conc}}{f_{dilution}}$$

At t>0:

(36) $$[IgG_{unbound}] = [IgG_{unbound}]_0 + x_t$$

(37) $$[\text{Fc}\gamma\text{Ia}] = [\text{Fc}\gamma\text{Ia}]_0 + x_t$$

(38) $$[IgG_{bound}] = [IgG_{bound}]_0 - x_t$$

$$K_D = \frac{k_{off}}{k_{on}}$$

Differential equation describing the dissociation:

(39) $$\frac{d[IgG_{bound}]}{dt} = -k_{on} * [\text{Fc}\gamma\text{Ia}] * [IgG_{unbound}] + k_{off} * [IgG_{bound}]$$

(40) $$\frac{d[IgG_{unbound}]}{dt} = \frac{d[\text{Fc}\gamma\text{Ia}]}{dt} = -\frac{d[IgG_{bound}]}{dt}$$

(41) $$\frac{dx}{dt} = k_{on}([\text{Fc}\gamma\text{Ia}]_0 + x_t) * ([IgG_{unbound}]_0 + x_t) - k_{off} * ([IgG_{bound}]_0 - x_t)$$

(42) $$\frac{counts(IgG_{bound})}{counts(IgG_{tot})} = \frac{[IgG_{bound}]_0 - x_t}{[IgG]_{tot}}$$
